# Integrating top-down and bottom-up approaches to understand the genetic architecture of speciation across a monkeyflower hybrid zone

**DOI:** 10.1101/2022.01.28.478139

**Authors:** Sean Stankowski, Madeline A. Chase, Hanna McIntosh, Matthew A. Streisfeld

## Abstract

Understanding the phenotypic and genetic architecture of reproductive isolation is a longstanding goal of speciation research. In many systems, candidate barrier traits and loci have been identified, but causal connections between them are rarely made. In this study, we combine ‘top-down’ and ‘bottom-up’ approaches with demographic modeling toward an integrated understanding of speciation across a monkeyflower hybrid zone. Previous work in this system suggests that pollinator-mediated reproductive isolation is a primary barrier to gene flow between two divergent red- and yellow-flowered ecotypes of *Mimulus aurantiacus*. Several candidate floral traits contributing to pollinator isolation have been identified, including a difference in flower color, which is caused primarily by a single large-effect locus (*MaMyb2*). Other anonymous SNP loci, potentially contributing to pollinator isolation, also have been identified, but their causal relationships remain untested. Here, we performed demographic analyses, which indicate that this hybrid zone formed by secondary contact, but that subsequent gene flow was restricted in a large fraction of the genome by barrier loci. Using a cline-based genome scan (our bottom-up approach), we demonstrate that candidate barrier loci are broadly distributed across the genome, rather than mapping to one or a few ‘islands of speciation.’ A QTL analysis (our top-down approach) revealed most floral traits are highly polygenic, with little evidence that QTL co-localize, indicating that most traits are largely genetically independent. Finally, we find little convincing evidence for the overlap of QTL and candidate barrier loci, suggesting that some loci contribute to other forms of reproductive isolation. Our findings highlight the challenges of understanding the genetic architecture of reproductive isolation and reveal that barriers to gene flow aside from pollinator isolation may play an important role in this system.

## Introduction

Understanding the phenotypic and genetic architecture of reproductive isolation is a major goal of modern speciation research [1–3]. Early studies took a ‘top-down’ approach by using quantitative trait locus (QTL) mapping and other association methods to detect genomic regions controlling barrier phenotypes or genetic incompatibilities [4–6]. More recently, ‘bottom-up’ approaches, such as genome scans of genomic differentiation (e.g., *F_ST_*) or admixture (e.g., *f_d_*), have identified candidate barrier loci in numerous systems, including those where isolation is thought to result from ecologically-based divergent selection or intrinsic incompatibilities [7–10].

Although both approaches have clear strengths, they also present significant challenges [11]. Top-down methods require that traits involved in reproductive isolation have already been identified, so our understanding of the genetic architecture of speciation can only ever be as complete as our knowledge of the traits controlling reproductive isolation in the system. In contrast, bottom-up approaches can provide a comprehensive view of the genomic landscape of speciation without complete knowledge of the isolating traits (but see [3, 12]). However, even though candidate barrier loci can be identified, their causal relationship with previously identified barrier traits usually remains unclear. This is because speciation usually involves many different isolating barriers (e.g., pre- and post-zygotic, extrinsic and intrinsic) [13, 14] that can become coupled together through different aspects of the speciation process [15–17]. Although the coupling of different barriers eases speciation by generating a stronger overall barrier [16, 17], the resulting linkage disequilibrium (LD) among barrier loci makes it difficult to understand their individual contributions to barrier traits. For example, a barrier locus identified in a genome scan might underlie an obvious phenotypic difference, or it may underlie a completely different barrier that is less conspicuous or that has yet to be discovered.

Therefore, instead of relying on one approach, many researchers have advocated for the integration of top-down and bottom-up methods [3, 11, 18]. However, this kind of integration is missing from most studies of speciation, meaning that any links between candidate barrier traits and barrier loci remain tentative. To date, some of the best efforts to integrate top-down and bottom-up analyses have made use of natural hybrid zones between divergent populations [19]. Hybrid zones have been described as natural laboratories, because they allow us to understand how reproductive isolation and barriers to gene flow play out in the real world [20]. In addition, their presence provides compelling evidence for ongoing gene flow between the taxa being studied, the relative duration of which can now be estimated using demographic inference methods. Moreover, cline theory provides a rich, spatially explicit framework for studying selection and gene flow across porous species boundaries [21–23]. Specifically, the shape and position of geographic clines are impacted by the relative effects of selection and gene flow across a hybrid zone [22]. Cline analysis has clear advantages over population genetic summary statistics used in most selection scans (e.g., *F*_ST_), and it can be applied to phenotypic traits [24].

However, it is only beginning to be applied to genome-scale datasets [24–28]. In this study, we combine top-down and bottom-up analyses to investigate the phenotypic and genetic architecture of pollinator isolation between hybridizing ecotypes of the bush monkeyflower (*Mimulus aurantiacus*).

In San Diego County, California, there is a sharp geographic transition between red- and yellow-flowered ecotypes of *M. aurantiacus* ssp. *puniceus* [29]. Despite being very closely related (*d_a_* = 0.005; [30], the ecotypes show extensive divergence across a suite of floral traits, including color, size, shape, and placement of reproductive parts (Fig. 1; [29, 31, 32]). Previous work suggests an important role of pollinators in driving floral trait divergence and reproductive isolation in this system [29, 33–36]. Field experiments have shown that hummingbirds and hawkmoths show strong preferences and constancy for the flowers of the red and yellow ecotypes, respectively [34, 36]. In addition to providing a source of divergent selection, pollinator behavior generates substantial premating isolation, potentially reducing gene flow between the ecotypes by 78% in sympatry [36]. Post-mating isolation is weak between the ecotypes, suggesting that pollinator isolation is the primary barrier to gene flow in this system [36].

**Figure 1.**
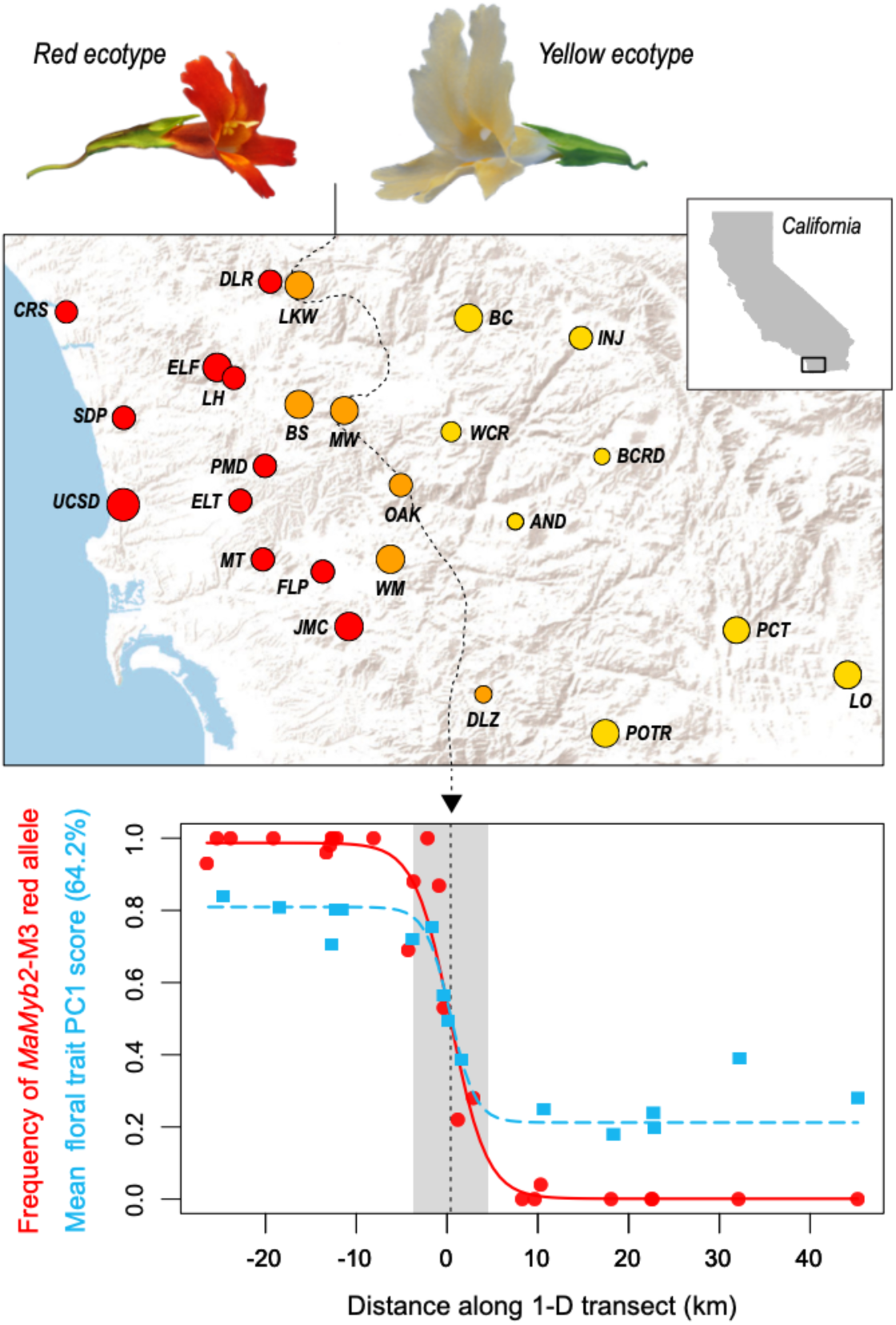
Clinal variation across a bush monkeyflower hybrid zone. (**top**) Typical flower phenotypes of the red and yellow ecotypes, and a map of the 25 sampling locations in San Diego County. The size of the circles shows variation in the sample sizes, which range from 4 to 18 individuals, totaling 292 individuals. The dashed line indicates the center of the hybrid zone, previously inferred from spatial variation in the frequency of alternative alleles at the M*aMyb2* locus. (**bottom**) Clines in allele frequency at the *MaMyb2* locus (red circles) and the mean floral trait PC1 score (blue squares) across the one-dimensional transect. The solid and dashed lines are the ML sigmoid cline models for *MaMyb2* allele frequency and trait PC1 score, respectively. The gray shaded rectangle represents the width of the hybrid zone.

**Figure 2.**
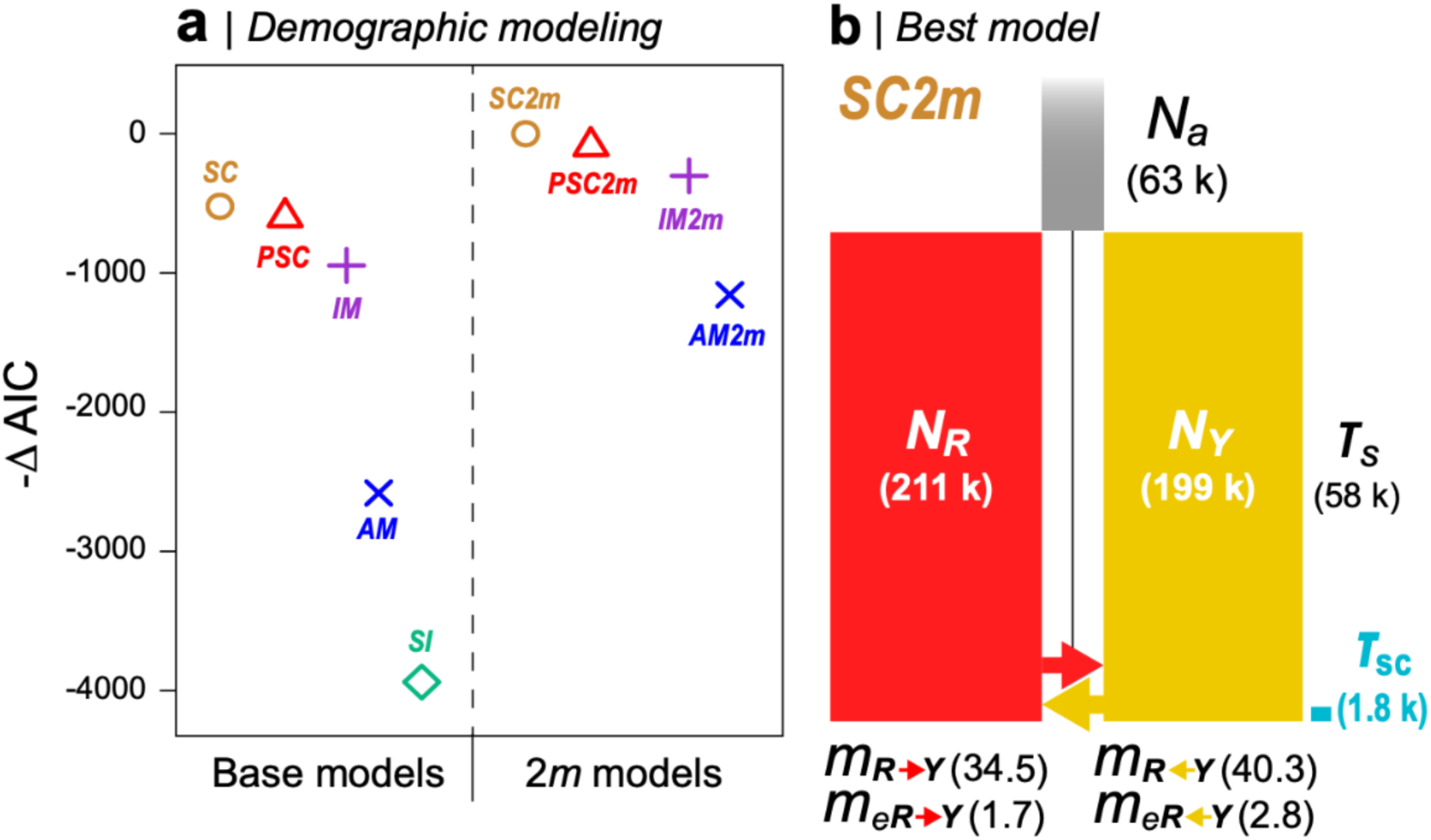
Demographic modeling reveals a history of gene flow following isolation. **(a)** - ΔAIC scores for the 9 demographic models fitted to the observed JSFS using *∂a∂i*. The base models (left of the dashed line) include a single migration parameter (*m*) for all loci, whereas the 2*m* models include separate migration parameters for neutral loci (*m*) and loci affected by a barrier to gene flow (*m*_e_). The best model (SC2*m*) has a -ΔAIC of 0, with more negative values indicating that models were a poorer fit. **(b)** A graphical depiction of the SC2*m* model. The width of the columns is proportional to the population size estimates for the ancestral (*N_a_*), red (*N_R_*), and yellow (*N_Y_*) populations. the height of the red and yellow bars is proportional to the total time in generations (*T_S_*) that has passed since the split. The blue bar shows the period during which secondary gene flow (*T*_SC_) occurred. The difference in arrow size is proportional to the difference in the bi-directional migration rate, *m*. The rates of effective migration (*m*e) are too small to show graphically.

Although the strength of pollinator-mediated reproductive isolation is strong, it is incomplete, meaning that there is potential for gene flow between the ecotypes in locations where their distributions overlap. This has led to the formation of a narrow hybrid zone, characterized by extensive phenotypic variation and geographic clines in several floral traits. For example, there is a steep cline in flower color that is centered on the hybrid zone and matches a similarly steep cline in the gene *MaMyb2*, which controls much of the variation in pigmentation [35]. Other floral traits and anonymous single-nucleotide polymorphisms (SNPs) also show clinal variation, implying that multiple traits contribute to reproductive isolation [24, 33]. However, evidence for an association between these phenotypic traits and genotypic signatures of selection is currently lacking.

In this study, we use demographic modeling, a cline-based genome scan, and QTL mapping to investigate the history of divergence and the connection between phenotypic and genomic signatures of selection in this system. One possible outcome is that QTL for the divergent phenotypes will overlap with regions of the genome under selection, as predicted if pollinator-mediated selection is the main barrier to gene flow between the ecotypes. These regions may be abundant and widespread across the genome, reflecting polygenic divergence, or they may consist of one or a few genomic regions enriched for loci that underlie multiple floral traits. Under an alternate scenario, we may find that floral QTL only partially overlap with genomic signatures of selection, which might reflect the spatial coupling of multiple different kinds of barriers. Our findings highlight the challenges of understanding the genetic architecture of reproductive isolation, and suggest that barriers to gene flow aside from pollinator isolation may also contribute to speciation in this system.

## Methods

### RAD sequencing, read filtering, and SNP calling

We identified SNPs using previously sequenced restriction-site associated DNA sequences (RADseq) from 292 individuals sampled from 25 locations across the hybrid zone (mean individuals per site = 12; range 4–18) [24]. These included 11 sites in the range of the red ecotype, 8 sites in the range of the yellow ecotype, and 6 sites in the hybrid zone (Table S1).

We processed the raw sequences, identified SNPs, and called genotypes using *Stacks* v. 1.41 [37]. Reads were filtered based on quality, and errors in the barcode sequence or RAD site were corrected using the *process_radtags* script in *Stacks*. Individual reads were aligned to the *M. aurantiacus* genome [30] using *Bowtie 2* [38], with the *very_sensitive* settings. We identified SNPs using the *ref_map.pl* function of *Stacks*, with two identical reads required to create a stack and two mismatches allowed when processing the catalog. SNP identification and genotype calls were conducted using the maximum-likelihood model (alpha = 0.01) [39]. To include a SNP in the final dataset, we required it to be present in at least 70% of all individuals; this resulted in a final dataset of 219,152 SNPs.

### Demographic inference

To gain a deeper understanding of the history of gene flow and selection in this system, we performed demographic inference in ∂a∂i [40]. We calculated the unfolded joint site frequency spectrum (JSFS) based on 19,902 SNPs, using subspecies *grandiflorus* as an outgroup to polarize alleles as ancestral or derived [41]. SNPs were included if they were genotyped in *grandiflorus* and in at least 70% of the red and yellow individuals. We included 124 individuals from 10 sites of the red-flowered ecotype and 65 individuals from 7 sites of the yellow-flowered ecotype, excluding sample sites that showed evidence of recent admixture (all hybrid sample sites and populations DLR and BC). The JSFS was projected to a sample size of 85 to maximize the number of segregating sites.

We fit nine two-population demographic models to the JSFS (Fig. S1): (*i*) strict isolation (SI), (*ii*) ancient migration (AM), (*iii*) isolation with migration (IM), (*iv*) secondary contact (SC), and (*v*) periods of secondary contact (PSC). The remaining four models—(*vi*) AM2*m*, (*vii*) IM2*m*, (*viii*) SC2*m*, and (*ix*) PSC2*m*—are the same as models *ii*-*v*, except that migration rates are inferred for two groups of loci to simulate the effect of a porous barrier to gene flow. For each model, we performed 20 independent runs using randomly generated starting parameters, and we reported the results for the run with the lowest log-likelihood. The goodness of fit of the models was determined using the AIC. Parameter estimates were converted into biologically meaningful values as described in [42], assuming a mutation rate of 7 x 10^-9^ [43].

### Admixture analysis

We used the model-based clustering program *Admixture* [44] to characterize patterns of genetic structure across the hybrid zone. We assigned the 292 individuals sampled from across San Diego County into two clusters (*K*=2) based on the full dataset of 219,152 SNPs that met our filtering requirements (note that *K*=2 was determined as the optimum number of clusters in [24]. In addition to using the full dataset, we also pruned SNPs using the *--indep-pairwise* function in *Plink* [45] to reduce linkage disequilibrium (LD) between neighboring SNPs (*r*^2^ threshold of 0.1, window size = 50 SNPs, step size = 10 SNPs).

We also ran *Admixture* separately for each chromosome, and for 2,173 non-overlapping windows, each containing 100 SNPs (mean window size of 89.1 kb with 8 - 38 RAD tags per window Fig. S2) The window-based analysis was automated using custom python scripts to produce plink.map and .ped files for each consecutive window, which were then passed to *Admixture*.

### Cline fitting

To quantify the geographic variation in ancestry (*Q*), we fit a sigmoid cline model to the mean ancestry scores from each site along a one dimensional transect, which was described in [24] (Fig. 1). Clines were fitted using *HZAR* [46] using the quantitative trait model, with the variance in the trait modeled separately on the left side, center, and right side of the cline. We estimated the following parameters: the cline center (*c*), defined as the inflection point of the sigmoid function; *Q*_left_ and *Q*_right_, the mean ancestry scores on the left and right sides of the cline, respectively; and the cline width (*w*), defined as the ratio between the total change in ancestry across the cline (Δ*Q*) and the slope at the cline center (note that Δ*Q* = *Q*_left_ - *Q*_right_ because we ensured that the mean ancestry score was higher on the left side before fitting). We conducted 3 independent fits with random starting values and retained the one with the highest log-likelihood. All of the best fits were visually inspected to ensure a sensible fit.

### Summarizing clinal variation in windows

After cline fitting, we calculated an *ad-hoc* statistic to identify genomic windows that had clines with a similar shape and position to the genome-wide cline. We refer to this statistic as the cline similarity score (*cs* score). Unlike individual parameters (*e.g.* the width or center), which describe a single feature of a cline, the *cs* score describes the shape and relative position of a cline with a single number. We calculate the cline similarity score as: *cs =* [Δ*Q*/(*W* + *l*)]*e*^(-(|*c*/*l*|))^2^^. Briefly (but see Supplement S1 for more details), the total change in ancestry, Δ*Q*, is divided by the sum of *w* and a scaling variable (*w* + *l*) to give an estimate of cline shape. The scaling variable controls the spread of shape scores across the joint distribution of Δ*Q* and *w*. In our case, *l* = half the length of the transect (0.5*t*), which results in high shape scores when clines have high Δ*Q* and low *w*, but low shape scores when clines have low Δ*Q* and high *w*. The shape score is then scaled according to the position of the cline center, *c*, relative to a position of interest. This could be a feature of the environment or a cline in a focal marker or trait. In our case, the position of interest is the center of the genome-wide ancestry cline. If the cline center coincides exactly with this point, then the shape score is equal to the *cs* score. However, the farther that the cline center is shifted away from the point of interest, the more the shape score is downgraded, resulting in a lower estimate of *cs*. Therefore, to have a high value of *cs*, a cline from a genomic window must have its shape and position closely match the cline in genome-wide ancestry (as in Fig. 3A). Finally, we scaled the *cs* score relative to the genome-wide ancestry cline, where 1 is the *cs* score calculated for the genome-wide ancestry cline, and 0 is the minimum value of *cs* observed for a window.

**Figure 3.**
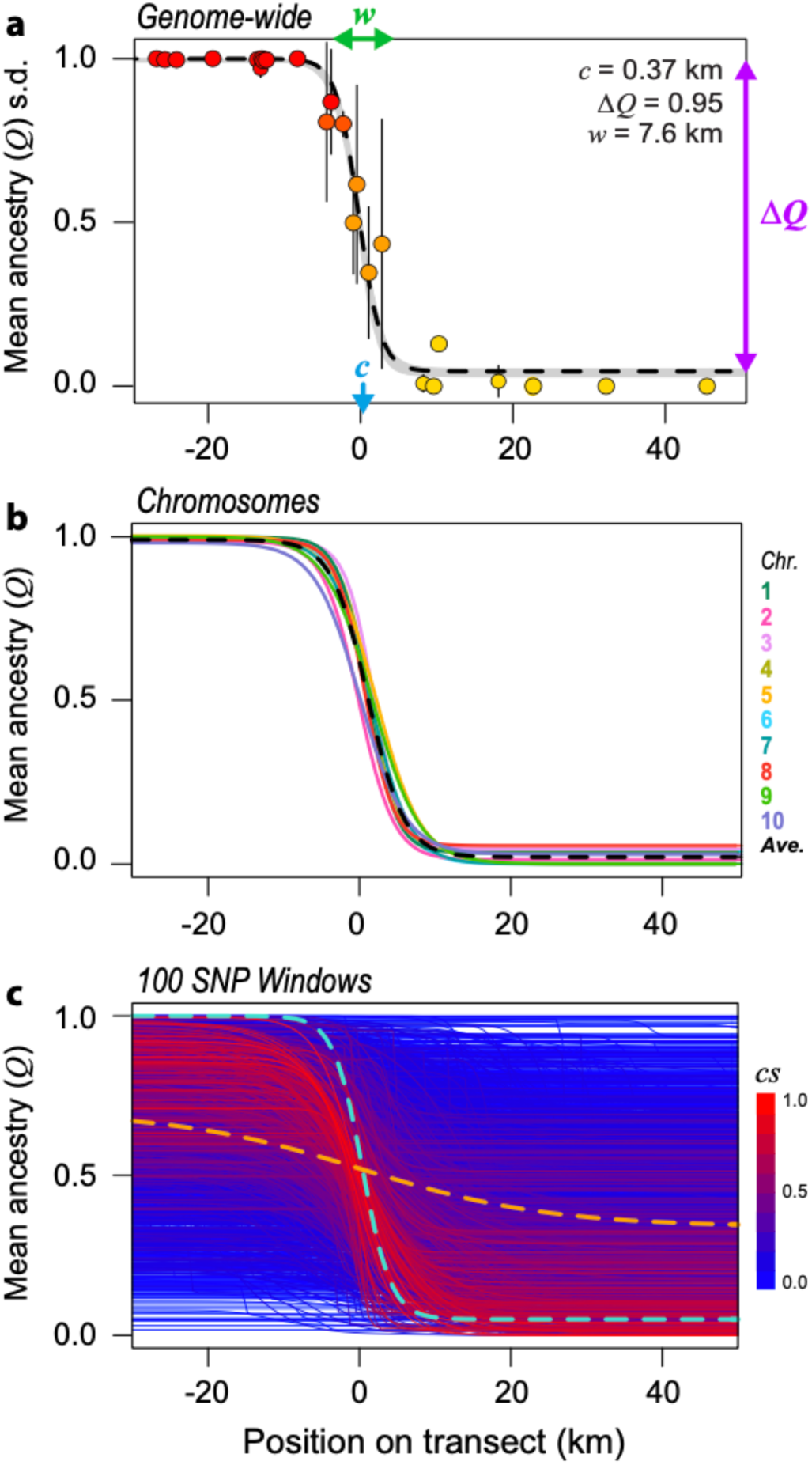
Clines in ancestry scores at different scales of genomic organization. **(a)** Genome-wide cline, inferred from the mean ancestry scores in each population along the 1-D transect. Position 0 on the horizontal axis corresponds to the cline center estimated from *MaMyb2* allele frequencies (see Fig 1b). The vertical bars show the standard deviation in ancestry scores for each population. The dashed line is the ML cline model, and the gray band is the two-unit support envelope. Three parameters of interest, including the cline center (*c*), width (*w*), and total change in ancestry across the cline (Δ*Q*), are indicated on the plot. **(b)** Ancestry clines estimated separately for each chromosome. Only the ML curves are shown for clarity (but see Fig S6). The dashed line is the mean cline, estimated by taking the average of the ML parameters for all chromosomes. **(c)** Ancestry clines estimated for 2,173, 100-SNP windows. The dashed cyan line shows the cline shape for the genome-wide cline (as shown in panel a), while the dashed orange line is the mean cline shape, estimated by taking the average of the ML parameters obtained for all windows. Each solid line is the ML sigmoid curve for one of the genomic windows. The curves are colored according to the value of the cline similarity score (cs), which indicates how similar the shape and position of each cline is to the genome-wide cline. Redder clines are more similar to the genome-wide cline and bluer clines are less similar (See main text for more details).

### Estimates of genetic differentiation in genomic windows

To characterize levels of genetic differentiation in a more traditional way, we calculated the population genetic statistic *F_CT_* between the ecotypes in each 100 SNP window using the program *Arlequin* [47]. This was done in an analysis of molecular variance framework that partitioned genetic variation between the ecotypes, among populations within ecotypes, and within populations. Populations were classified as coming either from the red or yellow ecotypes based on the *Admixture* results. Samples from hybrid populations were excluded from this analysis.

### QTL analysis

We used QTL analysis to identify genomic regions underlying divergent floral traits. We generated an outcrossed F_2_ population that contained 292 offspring produced by crossing two F_1_ individuals; each of these F_1_ parents was produced by crossing different greenhouse-raised red and yellow ecotype plants (from populations UCSD and LO; Table S1). To allow direct phenotypic comparison among plants grown in a common environment, we raised 25 red ecotype individuals (location UCSD), 31 yellow ecotype individuals (location LO) and 20 F_1_ individuals (LO x USCD) alongside the F_2_s. For each plant, we measured 13 floral traits (Fig. S3). Plants were raised as described in [33].

QTL mapping was conducted using *R/qtl* [48] and a previously published genetic map [30] generated from the same mapping population using *Lep-MAP2* [49]. We used phase information from *Lep-MAP2* to infer the grandparental origin of alleles in the F_2_s at 7574 mapped markers, which allowed us to recode them as coming either from a red or yellow grandparent. This set of markers was then reduced down to 2631—one per map position—by retaining the marker at each map position with the least missing data. Missing data for these markers was inferred by imputation using phase information from the mapping software. For each trait, we then used automated stepwise scanning for additive QTL and pairwise interactions. QTL identified using this procedure were then incorporated into a multi-QTL model to refine their positions, calculate 95% Bayes credible intervals, and estimate the percent phenotypic variation explained (i.e., the effect size) of each QTL.

### Test for an excess of QTL overlap

To test for co-localization among QTL, we used a permutation test to determine if there was significantly more overlap among QTL than expected by chance. We first estimated the observed number of overlaps based on the Bayes credible intervals among the 26 identified QTL using the *findOverlaps* function of the *GenomicRanges* package [50] in R and determined the average number of overlaps per QTL (*n* overlaps/*n* QTL). To determine if this statistic was significantly larger than expected by chance, we randomly generated new QTL positions while maintaining the observed number and size of observed QTL. We made the probability of QTL ‘landing’ on a given chromosome (Chr) a function of that chromosome’s length (*L_i_*) relative to the total genome length (*L*_T_), *P*(Chr*_i_*) = L*_i_* / L*_T_*, so that larger chromosomes were more likely to have QTL *i* assigned to them. We calculated the mean number of overlaps per QTL for 9,999 random datasets and estimated a *p*-value for the observed value as the number of permuted datasets where *n* overlaps/*n* QTL was equal to or greater than the observed estimate + 1/number of permutations +1.

### Test for overlap of QTL and outlier windows

We also used a permutation test to determine if QTL regions were enriched for outliers identified in our cline and *F*_CT_ based genome scans. We first counted the observed number of outlier windows within the empirical QTL intervals. This was performed for both *cs* and *F*_CT_ outliers, defined using two different cutoffs (top 1% and 5% of the empirical distributions). To determine if these counts were significantly different from chance, we produced 9,999 datasets where the genomic position of outlier windows was randomized, and we counted the number of outliers falling inside the empirical QTL intervals. A *p*-value for the estimate was calculated as described above.

## Results

### Evidence for a history of heterogeneous gene flow

The results of our demographic modeling support a history of gene flow across the range of the ecotypes. First, demographic models that included contemporary gene flow were far better at recreating the observed JSFS than a model of divergence without gene flow (i.e., strict isolation; -ΔAIC = 3938), or the best model of ancient migration, which included historical but not contemporary gene flow (-ΔAIC = 1157) (Fig. 2; Fig. S4). Second, models that included heterogeneous migration across the genome (2*m*) were always strongly favored over the equivalent models, where gene flow was modeled with a single rate (Fig. 2). Third, the SC and PSC models, which included periods of allopatry and secondary contact, were strongly favored over the IM model, where divergence occurred without a period of geographic isolation. The best-fitting model was the SC2*m* model, indicating that divergence of the red and yellow ecotypes included a period of allopatry followed by gene flow upon secondary contact (Fig. 2). Assuming a mutation rate of 7×10^-9^ [43], the ML parameters indicate that the ecotypes have been exchanging an average of 37 migrants per generation (*m_YR_* = 34.5 per gen.; *m_RY_* = 40.3 per gen.) for the last 1,800 generations, which equates to roughly 3,600 years, assuming a two-year generation time for these perennial plants. Despite evidence that gene flow between the ecotypes has been extensive, the ML model suggests that 37.4% of loci have experienced a substantial reduction in effective migration (15-20 fold; *m*_e*YR*_ = 1.7 per gen.; *m*_e*RY*_ = 2.7 per gen.) due to the effects of selection against gene flow (Fig. 2).

### Sharp clines in genome- and chromosome-wide ancestry

The presence of sharp clines in multiple floral traits suggests that some fraction of the genome is impacted by selection against gene flow [33]. The results from *Admixture* support these findings, revealing genome-wide patterns of ancestry that closely match the ecotypic designations assigned based on floral phenotypes (Fig. S5). Specifically, red- and yellow-flowered individuals sampled from either side of the hybrid zone were strongly assigned to alternate clusters, while individuals from hybrid populations tended to show some assignment to both clusters, indicating their genomes are a mix of red and yellow ancestry. The results are nearly identical between independent runs of *Admixture* that include the full dataset or subsets of the data pruned to minimize LD between neighboring SNPs (*r*^2^ > 0.999).

To compare these changes in ancestry to the observed geographic variation in floral traits, we used cline analysis to fit a sigmoid cline to the mean ancestry scores from each site. The best-fitting cline model provides an excellent summary of the change in ancestry across the transect (Fig. 3a) and has an extremely similar shape to cline models from the divergent floral traits and molecular markers (Fig. 1). In addition, consistent with the increased variance we observed in multiple phenotypic traits in hybrid populations [33], the standard deviation of ancestry scores is higher in sample sites close to the cline center, thus providing genomic evidence for hybridization (Fig. 3a).

The *Admixture* scores provide additional genetic evidence for restricted gene flow across the hybrid zone, but they give us no indication as to the number of loci involved or their genomic distribution. For example, the differences in ancestry could be driven by a small number of loci that reside on a single chromosome, or they could reflect more widespread genomic divergence, involving loci scattered across multiple chromosomes. By repeating the cline analysis of ancestry scores separately for each chromosome, we find highly consistent clines in ancestry for all 10 chromosomes (Fig. 3b, Fig. S6).

### Heterogeneous clinal variation across the genome

To understand how cline shape varies at a finer genomic scale, we fit clines to 2173 non-overlapping 100-SNP windows. This analysis revealed broad variation in geographic patterns of ancestry (Fig. 3c). Unlike each chromosome, the majority of windows show little or no spatial change in ancestry between the red and yellow ecotypes, translating into very low *cs* scores (mean *cs* = 0.15; s.d. = 0.15, Fig. 4).

**Figure 4.**
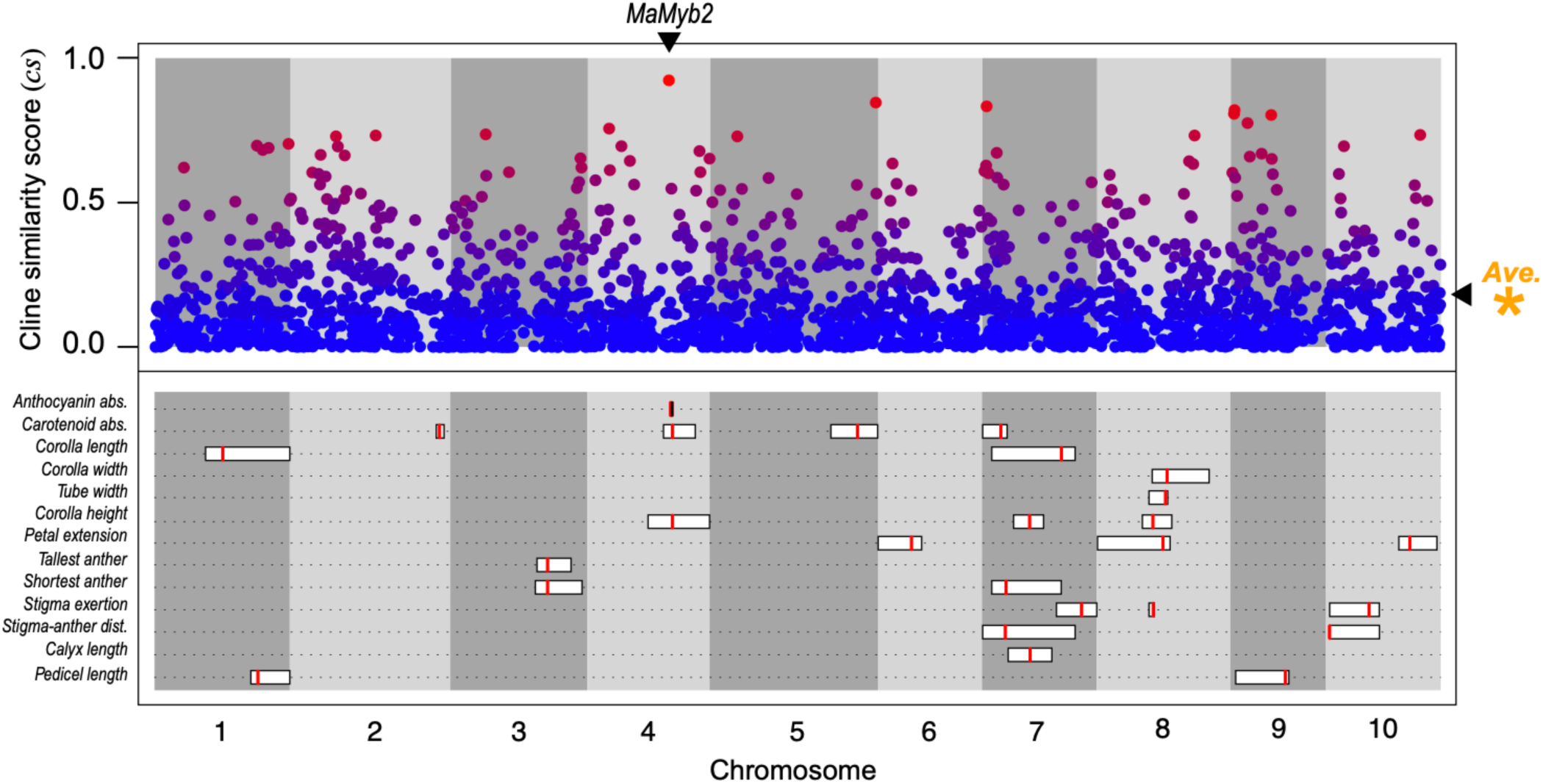
Cline-based genome scan and locations of QTL for floral traits. (top) The scaled cline similarity (*cs* score) score in each 100 SNP window plotted against the physical position of the window in the bush monkeyflower genome. The points are colored as in Figure 3c, with redder points containing windows with *cs* scores that are more similar to the genome-wide pattern and bluer points are less similar (See main text for more details). The orange asterisk denotes the average barrier score among all windows. The position of the *MaMyb2* gene that controls differences in flower color is shown. **(bottom)** The positions of the QTL for the 13 measured floral traits plotted along the physical position of the genome. The red vertical line corresponds to the best estimate of the QTL peak, and the width of the rectangles denotes the 95% Bayes credible intervals of the estimated QTL position.

However, some genomic regions show clines in ancestry that strongly resemble the genome wide cline, suggesting that they contain barrier loci. This includes the window with the highest *cs* score (*cs =* 0.91), which contains the known barrier locus, *MaMyb2*. The shape and position of this window-based ancestry cline (Δ*Q* = 0.95, *w* = 10.4 km, *c* = -0.3 km) is highly similar to the genome-wide cline in ancestry (Δ*Q* = 0.95, *w* = 7.6 km, *c* = 0.37 km) and to the cline in allele frequency for a SNP in *MaMyb2* (*MaMyb2*-M3 marker: Δ*P* = 0.99, *w* = 8.1 km, *c* = -0.07 km [24]. However, rather than a clear set of outliers, we observe a continuous distribution of *cs* scores. Therefore, we use two arbitrary cutoffs (top 1% and 5% of the distribution of *cs* scores) to define a set of candidate windows potentially containing barrier loci.

Regardless of which cutoff we use, these candidate barrier regions are broadly distributed across the genome. For the 5% cutoff, they occurred on all 10 chromosomes (5 - 20 windows per chromosome; for the 1% cutoff, they occur on 9 of the 10 chromosomes with 1 - 4 windows per chromosome). There were only 12 cases where candidate windows were directly adjacent, indicating that they also were broadly distributed within each chromosome. We also find that genetic differentiation is higher for candidate regions than for the genomic background (1% mean *F_CT_ =* 0.31, 5% mean *F_CT_ =* 0.23, overall mean *F_CT_* = 0.07). However, *F_CT_* explains only 38% of the variation in *cs* scores (Fig. S7).

### Most candidate barrier traits are polygenic

We used quantitative trait locus (QTL) mapping to identify regions of the genome associated with candidate barrier traits. The 13 floral traits showed significant differences between pure red and yellow ecotype plants when grown in a common environment, with mean trait values differing by 0.9 to 7.1 standard deviations (Fig. S8). A total of 26 QTL were identified. For nine traits, we identified more than one QTL (range 2 – 4), and QTL were located on all 10 linkage groups, with LG 7 containing QTL for seven different traits (Fig. 4, Figs. S9 and S10). On average, each QTL explained 9.9% of the variation in the F_2_ population (range 1.82% - 62.6%) (Fig. S11), with an average total variation explained for each trait of 19.8%. The exception was a large-effect QTL for anthocyanin content on LG 4 that explained 62.6% of the variation and mapped to a region near the previously identified causal locus *MaMyb2* [35]. Thus, despite clear heritable differences in these traits, QTL analysis was able to explain only a modest amount of the segregating variation, indicating that most traits have a polygenic architecture.

The presence of multiple QTL occurring on the same chromosome indicates that some regions may contribute to multiple traits, which would help maintain trait associations in hybrid offspring [17]. Overall, we find that QTL do tend to co-localize more often than would be expected by chance (mean observed overlap of 3.23 QTL; mean permuted overlap = 2.51 QTL; *p* = 0.042, Fig. S12). However, the effects of this co-localization are seen most strongly only for size-related traits (e.g. corolla length and height of the tallest anther), which remain highly correlated in the F_2_ generation (*r* = 0.88) (Fig. S13). By contrast, the average correlation coefficient among all other pairs of traits was much lower (mean absolute value of *r* = 0.26). For example, the three overlapping QTL on LG 4 that control anthocyanin and carotenoid pigmentation, as well as corolla height, span a total physical distance of only 76 kb. However, these traits show weak correlations in the F_2_ population (anthocyanin *vs* carotenoid: *r* = 0.18; anthocyanin *vs* corolla height: *r* = -0.20; carotenoid *vs* corolla height: *r* = 0.19). This shows that the QTL overlap would have little effect on maintaining the phenotypic correlations where hybridization occurs.

### Low concordance between QTL and outlier regions

Finally, we tested for overlap between the floral trait QTL and the candidate barrier regions from the cline-based and *F_CT_* genome scans. Using a permutation test, we tested whether genomic windows with higher *cs* scores tended to overlap with QTL more often than expected by chance. Regardless of which cutoff we used (e.g. top 1% or top 5% of *cs* scores), we found that floral trait QTL were not significantly enriched for candidate barrier regions (p > 0.3, Fig. S14). This suggests a complex connection between the genetic and phenotypic architecture of reproductive isolation. The results were the same when we defined candidate barrier regions based on *F_CT_* (e.g. top 1% or top 5% of the *F_CT_* distribution; Fig. S14).

Given that wide QTL intervals reduce the power of the enrichment test, we also asked how often the estimate of the QTL peak fell within a candidate barrier window. However, even when using the 5% cutoff, we found that none of the QTL peaks occurred within candidate barrier regions. This included the QTL for floral anthocyanin, where the QTL peak occurs 589 kb from the window containing the causal locus, *MaMyb2*.

## Discussion

In this study, we used a combination of QTL mapping and population genomic analyses to obtain a deeper understanding of the phenotypic and genetic architecture of reproductive isolation in a hybrid zone. Past studies in this system have separately identified candidate floral traits contributing to pollinator-mediated reproductive isolation [33, 34], and anonymous, candidate barrier loci [24]. Here, we use top-down and bottom-up approaches in an effort to connect phenotypic and genetic candidates. These results are discussed in the context of new insights about the history of divergence revealed by demographic analysis and are aided by a known, large-effect barrier locus (*MaMyb2*) with a clear phenotypic effect.

### The history of divergence: new insights from demographic analysis

A firm understanding of the historical demography of speciation is essential when interpreting divergence across hybrid zones [51, 52]. In zones that are at demographic equilibrium, it is possible to interpret clines in terms of migration, selection, and drift and sampling effects [22]. However, if hybrid zones formed recently, clines in neutral loci or traits will be steep initially, but they will decay over time [22]. Previous work in this system revealed patterns of isolation-by-distance across (and orthogonal) to the hybrid zone that were consistent with a long-term ‘stepping-stone’ pattern of gene flow across the entire range of the study area [33]. In fact, there was no evidence for substantial genome-wide differentiation between the ecotypes after correcting for the effect of geography. Based on this result, Stankowski *et al*. [33] concluded that the hybrid zone formed due to one of three possible scenarios: (i) a primary origin with continuous gene flow during divergence, (ii) a secondary origin, where divergence occurred in allopatry, followed by extensive gene flow after contact resumed, or (iii) a secondary origin where the period of allopatry was short.

Our demographic analyses provide new evidence that this hybrid zone formed from secondary contact after a long period of isolation. Indeed, both models that included periods of geographic isolation (SC and PSC) were a better fit to the data than models of continuous gene flow (IM). The parameter estimates for the preferred model (SC2*m*) indicate a relatively long period of isolation, followed by a period of secondary contact that began roughly 1,800 generations ago. It is important to note that we fit relatively simple models to the data that excluded changes in population size in the ancestral and daughter populations and variation in *N_e_* along the genome. Recent work has shown that failure to model key parameters can result in incorrect inference under some circumstances [53]. Although more sophisticated modelling may arrive at different conclusions in the future, these results point to a secondary origin of this hybrid zone.

In terms of the main goal of our paper, the key result of the demographic analysis was that all models with two rates of migration (2*m* models) fit the data better than those where migration was modelled at a single rate, indicating a heterogeneous pattern of gene flow across the genome. Moreover, the estimated parameters for the preferred model suggest that roughly one third of loci (37%) have experienced migration at substantially reduced rate compared with non-barrier loci. This result supports previous conclusions that candidate barrier traits and loci are indeed impacted by natural selection [24], further motivating the need to connect the phenotypic and genetic architecture of RI in more detail.

### From the ‘bottom up’: insights from the cline-based genome scan

Starting with the bottom-up approach, we used a cline-based genome scan to identify candidate loci underlying barriers to gene flow. Unlike traditional summary statistics calculated between pre-defined groups (e.g., *F_ST_*), geographic cline analysis is a more natural framework for studying genetic divergence across hybrid zones, as it provides clearer insights into the strength and nature of selection [24, 26]. Rather than fitting clines to allele frequencies for individuals SNPs, we fit clines to model-based ancestry scores, treating them as a quantitative trait [21].

By conducting this analysis at different scales of genomic organization, we are able to conclude that candidate barrier regions are widespread throughout the genome. At the genome-wide scale, the cline in ancestry is centered on the hybrid zone and has a very similar shape to the clines in floral traits. At the chromosome scale, all 10 chromosomes show clear sigmoid clines in ancestry, with their shapes and positions being highly similar to the genome-wide cline. The same conclusion can be drawn from the window-based analysis, as candidate barrier regions are present on all 10 of the chromosomes. The window with the highest *cs* score is located on chromosome 4 and contains the gene *MaMyb2*. Alternate alleles at this locus control the difference in flower color, and other population genetic analyses indicate that it has been subject to strong divergent selection in this system [35, 54]. Prior knowledge of this barrier locus provides confidence that other windows with high *cs* scores also likely harbor barrier loci with similarly large phenotypic effects.

However, rather than identifying a clear set of *cs* outliers, we observed a continuous distribution of *cs* scores, indicating that clines show varying degrees of resemblance to the genome-wide cline. Although it is tempting to interpret variation in the cline similarity score exclusively in terms of the sieving effect of a porous species boundary (i.e., assuming that the *cs* score is proportional to a local reduction in gene flow caused by associated barrier loci), the observed variation in *cs* scores requires more conservative interpretation. First, neutral processes, such as isolation-by-distance, can generate clines that are highly similar to clines generated by selection [26]. Similarly, neutral clines generated by secondary contact can take a long time to decay, making them hard to distinguish from selected ones [22, 51]. Localized drift (and sampling effects) tends to distort cline shapes in a way that may lead to the discovery of false positives [55, 56]. In addition, even though a genomic region may contain a large-effect barrier locus, it might not show a cline if the genomic window is too broad to capture the relevant signatures. Future efforts to help identify non-neutral clines may be accomplished using whole-genome rather than reduced-representation sequencing, and by comparing results obtained from multiple hybrid zones [27]. Simulations of cline formation could also help distinguish candidate outliers (as in [26]). Even with these measures in place, the noise generated by background processes and sampling effects may mean that we only have power to confidently detect large-effect loci, which remains a general problem with all genome scan approaches [3].

### From the ‘top down’: insights from QTL analysis of candidate isolating traits

We mapped QTL to identify genomic regions underlying floral trait divergence in this system. Although we identified a small number of QTL for each trait (between 1 and 4), the identified QTL explained only about 20% of the variation in each trait. Given that these traits are under strong genetic control, the ‘missing’ variance implies that most of the floral traits are polygenic, caused by many loci with effect sizes below our limit of detection. However, some fraction of the unexplained variation also may be due to subtle environmental differences experienced by each plant in the greenhouse.

Although finding many small-effect loci may be expected in studies of phenotypic evolution [57], many analyses of adaptation and speciation have recovered distributions of effect sizes skewed toward larger effects [58, 59]. Moreover, the identified regions often control more than one trait, and in some cases, more than one type of isolating barrier (e.g., pre- and post-mating barriers). For example, in another pair of *Mimulus* species, *M. cardinalis* and *M. lewisii*, large-effect QTL for multiple traits associated with pollinator isolation and hybrid sterility occur in a few genomic regions thought to harbor chromosomal inversions [4, 60]. In sunflowers (*Helianthus*), multiple traits are associated with local adaptation to dune and non-dune habitats and map to a small number of large, non-recombining haplotypes containing structural variants [61] (see [62–64] for other examples).

Large, multiple effect loci are an expected outcome of local adaptation and speciation [17, 65], because more ‘concentrated’ genetic architectures are favored in scenarios where gene flow opposes adaptive divergence [65]. Not only do large effect loci make individual traits more visible to selection, but tight linkage and pleiotropy enhance the coupling of different sets of adaptive traits, meaning that they can remain associated despite gene flow [17]. This begs the question: why do we see so few large effect loci and such little overlap among floral trait QTL in the red and yellow ecotypes? One potential explanation is that divergence was initiated during a period of geographic isolation—a hypothesis that is supported by our demographic analysis. If trait divergence did occur during a phase of allopatry, the selection favoring certain combinations of traits could build-up LD among many small-effect loci without opposition by gene flow.

Although the associations would decay rapidly upon secondary contact (as we see in the hybrid zone; [24, 33], this decay would be expected to occur over a spatial scale determined by the strength of selection and the migration rate. If two adjacent habitats occur over a scale that is many times larger than the dispersal distance of the organism (as is the case between the red and yellow ecotypes; [33]), then hybridization has almost no bearing on adaptation occurring in distant parts of the range [66]. This makes divergence in parapatry almost as easy as in allopatry [67], meaning that local adaptation will persist far from the hybrid zone, and strong associations among small effect loci can remain in all regions except for those closest to the hybrid zone.

Another factor that may have influenced the outcome of our QTL analysis is that the parents we used for mapping came from populations located very far from hybrid zone. Because concentrated genetic architectures evolve as a response to gene flow, theory predicts that the genetic architecture of local adaptation may vary spatially in a way that reflects the local hybridization risk. This was highlighted in a hybrid zone between two ecologically differentiated subspecies of *Boechera stricta* [68]. In the vicinity of the hybrid zone, several phenological traits map to a single locus containing a chromosomal inversion, where the ecotypes are differentially fixed for the standard and inverted arrangements. However, in areas that are more distant to the hybrid zone, both ecotypes harbor the standard arrangement, and QTL mapping with these populations revealed distinct QTL. Although it is possible that the genetic architecture of divergence also varies across the range of the red and yellow ecotypes in a way that might favor divergence, our observations suggest that this is unlikely. Specifically, there is no evidence for the phenotypic maintenance of two distinct ecotypic forms within the hybrid zone. Instead, we see a continuum of phenotypic variation that resembles what we observe in the F_2_ mapping population [33], suggesting a similarly complex genetic architecture across the range.

### Inferences from integrating top-down and bottom up approaches

Having identified a set of candidate barrier loci and QTL regions for putative barrier traits, we next sought to understand how they were connected in relation to previous hypotheses about reproductive isolation in this system [24]. Taken at face value, the two analyses seem consistent, as they both suggest that divergence in this system is polygenic, involving regions spread across the genome. However, when we intersect the regions identified by these approaches, we find very little concordance. What does this tell us about divergence in this system?

First, there are some technical and biological explanations that could account for these findings. The first is that QTL analysis often has low resolution. Specifically, the QTL intervals are very wide, substantially reducing our power to test if QTL regions are enriched for candidate barrier loci. We see the same result if we focus on the estimated location of the QTL peaks, which never fall inside a candidate barrier region. This is even true for the peak of the large-effect QTL for floral anthocyanin pigmentation that is located 589 kb from the window with the highest *cs* score and contains the *MaMyb2* gene, which we know is responsible for the major difference in flower color [35]. Therefore, had we not had *a priori* knowledge about the position of the gene from previous work, it is likely we would have failed to connect this strong signal of selection with the underlying gene involved. In addition, our genome scan is based on RADseq data, so the SNP density may have been too sparse to obtain sequences in LD with the loci under selection. Moreover, from a biological perspective, the QTL analysis implies that most of the traits studied are polygenic, meaning that selection on each locus is weak, making it difficult to detect them using any genome scan. All of these factors probably contribute to the highly complex pattern that we see.

Similarly, although many of the candidate barrier loci have clines that resemble the window that contains the large effect locus *MaMyb2*, these windows are not associated with large-effect QTL for floral traits. One possible explanation for this is that we may have failed to measure the relevant floral traits contributing to pollinator isolation. This seems unlikely, given how well floral morphology has been studied in this system [29, 31–33]. We therefore hypothesize that other barriers to gene flow, besides pollinator isolation, play an important role in the maintenance of this hybrid zone. For example, local adaptation of non-floral traits could be an important source of pre- and post-mating isolation [13, 69]. We previously identified clines in several eco-physiological traits that may be associated with habitat-based isolation [70], but these show very shallow linear gradients rather than sharp sigmoid clines, and they are well explained by variation in local climate variables, making them unlikely candidates [70].

However, other ecologically-based barriers may exist that remain to be characterized. Finally, it is possible that some of the candidate barrier regions contain intrinsic incompatibilities that cause reduced fitness in hybrids [71]. Although our previous work found little evidence for intrinsic postzygotic isolation in the F_1_ generation, we did detect partial male sterility in some inter-ecotype crosses [36]. In addition, we have only surveyed plants under benign greenhouse conditions and only through the F_1_ generation. Genetic incompatibilities in later generations [72] or under natural conditions [73] might play a larger role in the maintenance of the ecotypes than anticipated—a prediction that has been made in relation to ‘ecological speciation’ more broadly [15].

### Conclusions and implications for studying the architecture of speciation

By combining top-down and bottom-up approaches with demographic modelling, our study provides new insight into the history and genetic architecture of speciation between these monkeyflower ecotypes. Our demographic analysis suggests that the hybrid zone formed by secondary contact, but gene flow following contact is now heterogeneous across the genome due to the effect of multiple barrier loci. A cline-based genome scan indicates that candidate barrier loci are widespread across the genome, rather than being associated with one or a few ‘islands’ of speciation. A QTL analysis of floral traits identified many QTL of small effect, with limited co-localization among QTL for different traits. We found limited evidence that QTL and candidate barrier loci overlap, suggesting that other barriers to gene flow aside from pollinator isolation may contribute to speciation.

In addition to providing knowledge about this system, our study has important implications for efforts to understand the phenotypic and genetic architecture of isolating barriers. For many study systems, candidate barrier traits and loci are identified in separate studies, meaning that the link between them is not tested explicitly. However, any barrier locus associated with an ecological gradient may underlie a completely different type of barrier. This was highlighted by [15], who showed how intrinsic barriers can become spatially coupled with ecologically-based barriers—a phenomenon that may cause researchers to erroneously identify incompatibility loci as those underlying local adaptation. The same issue also arises if multiple ecological gradients change in concert. We therefore advocate for additional studies that integrate top-down and bottom-up approaches before drawing conclusions about causal associations between candidate barrier traits and loci. Finally, our study shows that, even with a concerted effort, understanding the phenotypic and genetic basis of speciation is extremely difficult. Although emerging methods and data may help, this will likely remain a major challenge for the field.

## Data availability

Sequence data are available on the short read archive (SRA) under bio projects PRJNA299226 and PRJNA317601. All other datafiles are available on dryad. Scripts for performing custom analyses are available at https://github.com/seanstankowski/Mimulus_cline_QTL_manuscript.

## Acknowledgments

We thank Julian Catchen for making modifications to *Stacks* to aid this project. Peter L. Ralph, Thomas Nelson and Roger K. Butlin and Anja M. Westram and Nicholas H. Barton provided advice, stimulating discussion, and critical feedback. The project was supported by National Science Foundation grant DEB-1258199.

## Author contributions

SS and MAS designed the study and conducted the molecular work. SS and HM performed the QTL analysis. SS, MAC, and MAS conducted all other analyses. SS and MAS wrote the manuscript, with input from the other authors. Scripts for performing custom analyses are available at https://github.com/seanstankowski/Mimulus_cline_QTL_manuscript.

## Supplementary tables and figure captions

**Table S1.** Location information and samples sizes for the sample sites used in the study. Population codes correspond with those in Figure 1.

| Pop name | Latitude | Longitude | 1-D distance<br>(in meters) | *N* |
| --- | --- | --- | --- | --- |
| CRS | 33.13037 | -117.30717 | -26461 | 11 |
| UCSD | 32.8894 | -117.23618 | -25369 | 12 |
| SDP | 32.9981 | -117.23538 | -23857 | 12 |
| MT | 32.82095 | -117.06175 | -19135 | 12 |
| JMC | 32.73732 | -116.9541 | -13303 | 18 |
| ELF | 33.08595 | -117.1453 | -12893 | 11 |
| LH | 32.80582 | -116.9867 | -12730 | 12 |
| FLP | 32.89422 | -117.08982 | -12557 | 12 |
| ELT | 33.06088 | -117.11877 | -12168 | 12 |
| PMD | 32.93787 | -117.05913 | -8097 | 12 |
| WM | 32.82133 | -116.90228 | -4298 | 17 |
| BS | 33.0148 | -117.01643 | -3710 | 18 |
| DLR | 33.16818 | -117.05237 | -2153 | 9 |
| MW | 33.00718 | -116.95978 | -827 | 17 |
| OAK | 32.91407 | -116.88932 | -363 | 12 |
| LKW | 33.16372 | -117.0161 | 1177 | 15 |
| DLZ | 32.6525 | -116.78597 | 2891 | 4 |
| WCR | 32.9805 | -116.826517 | 8277 | 12 |
| AND | 32.868233 | -116.7461 | 9643 | 4 |
| BC | 33.12262 | -116.80468 | 10311 | 13 |
| POTR | 32.6038 | -116.63392 | 18064 | 11 |
| INJ | 33.09785 | -116.66432 | 22509 | 11 |
| BCRD | 32.94958 | -116.63795 | 22615 | 6 |
| PCT | 32.73258 | -116.46983 | 32097 | 7 |
| LO | 32.6767 | -116.33123 | 45224 | 12 |

**Table S2.** Estimated parameters from each demographic model. N_a_, size of the ancestral population; *N_r_*, size of the red population, *N_y_*, size of the yellow population; *T_s_*, duration of the split; *m*_ry_, migration from yellow into red; *m*_yr_, migration from red into yellow; *T_am_*, duration of ancient migration; *T_SC_*, duration of secondary contact.

**Figure S1.**
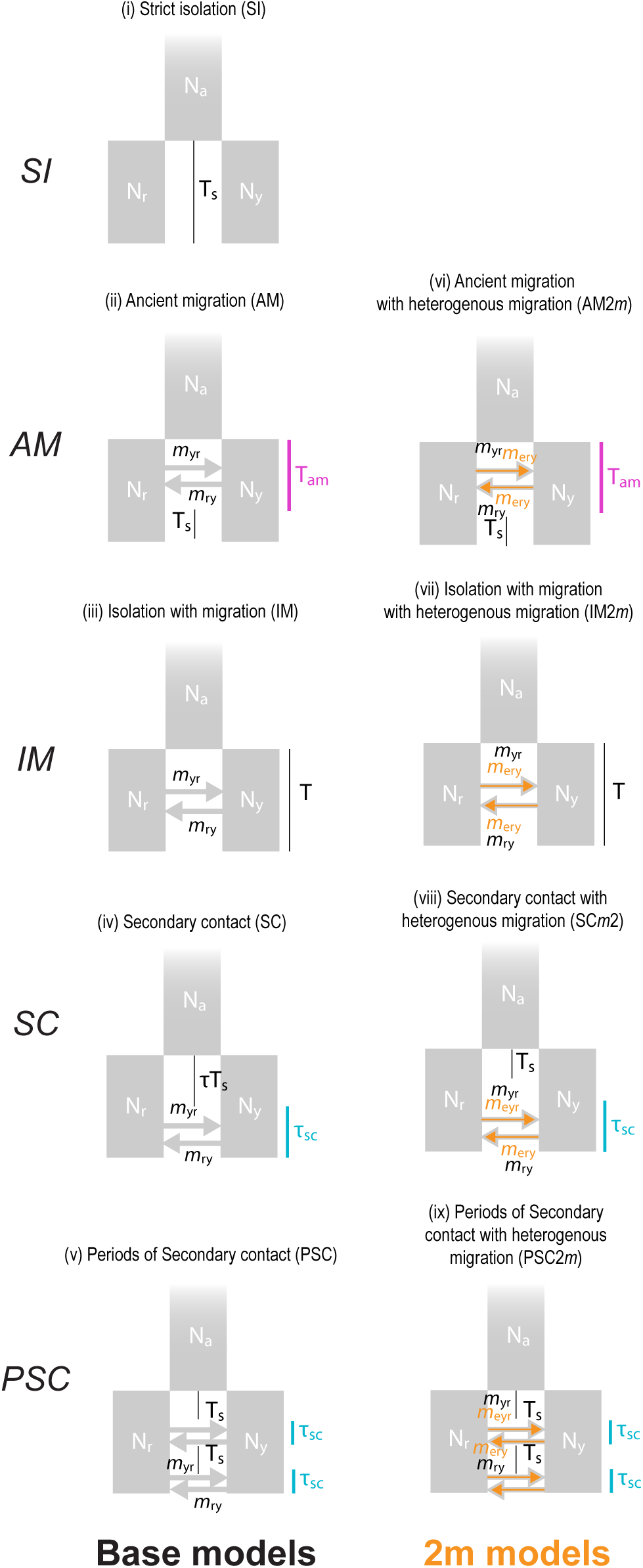
Cartoons of each model tested in the demographic analysis. (*i*) SI, strict isolation; (ii) AM, ancient migration; (*iii*) IM, isolation with migration; (*iv*) SC, Secondary contact; (*v*) PSC, Periods of secondary contact. The remaining four models—AM2*m,* IM2*m,* SC2*m*, PSC2*m*—are the same as models *ii*-*v*, except that migration rates are inferred for two groups of loci to simulate the effect of a porous barrier to gene flow. The model parameters are as follows: *N_a_*, size of the ancestral population; *N_r_*, size of the red population, *N_y_*, size of the yellow population; *T_s_*, duration of the split; *m*_ry_, migration from yellow into red; *m*_yr_, migration from red into yellow; *m_e_*_ry_, effective migration from yellow into red; *m_e_*_yr_, effective migration from red into yellow; *T_am_*, duration of ancient migration; T_SC_, duration of secondary contact.

**Figure S2.**
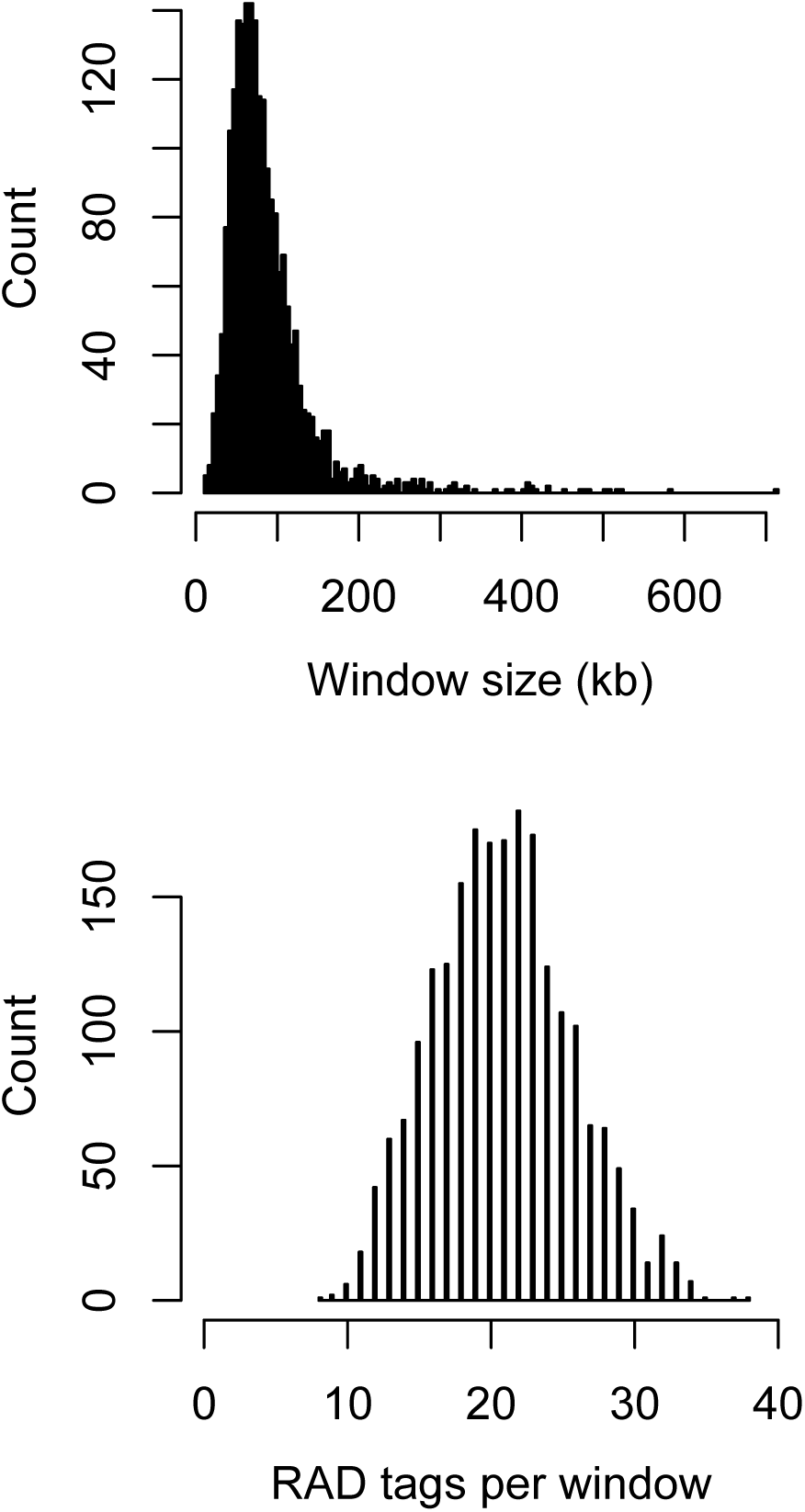
Information on the size and content of the 100 SNP windows. Distributions of window size (kb) and the number of RAD tags sequenced (post filtering) within each window.

**Figure S3.**
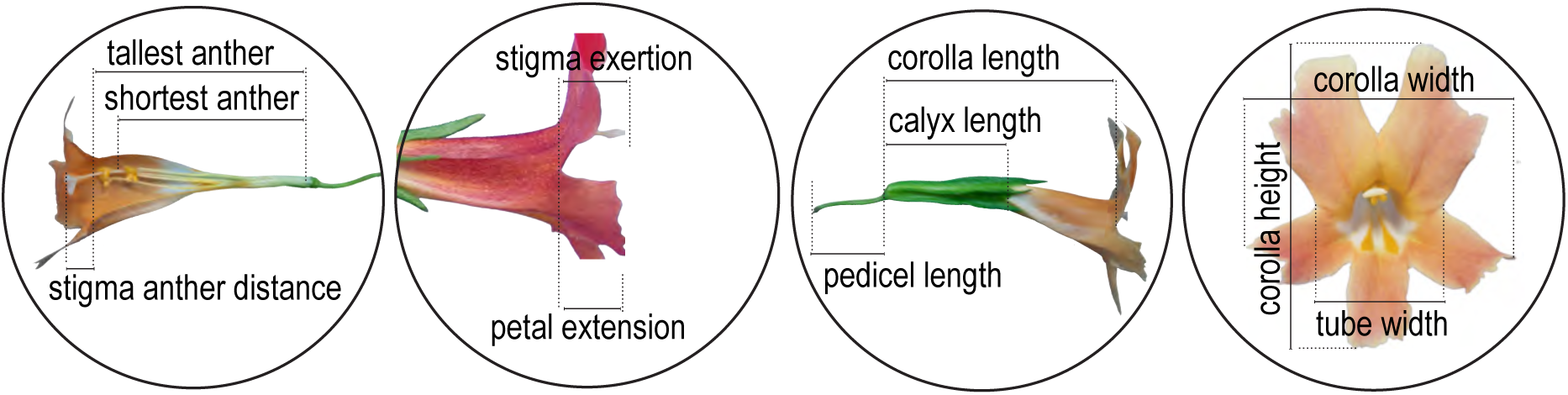
Floral traits measured for the QTL analysis. In addition to the size-related traits, we also measured anthocyanin and carotenoid pigment levels. Anthocyanins were extracted in 1% acidic methanol from a single disc collected from one of the top petals the first day each flower opened. Absorbance of extracts was measured with a spectrophotometer at 520 nm, as described previously [74]. Carotenoids were extracted in hexane following a similar protocol, and absorbance was measured at 450 nm.

**Figure S4.**
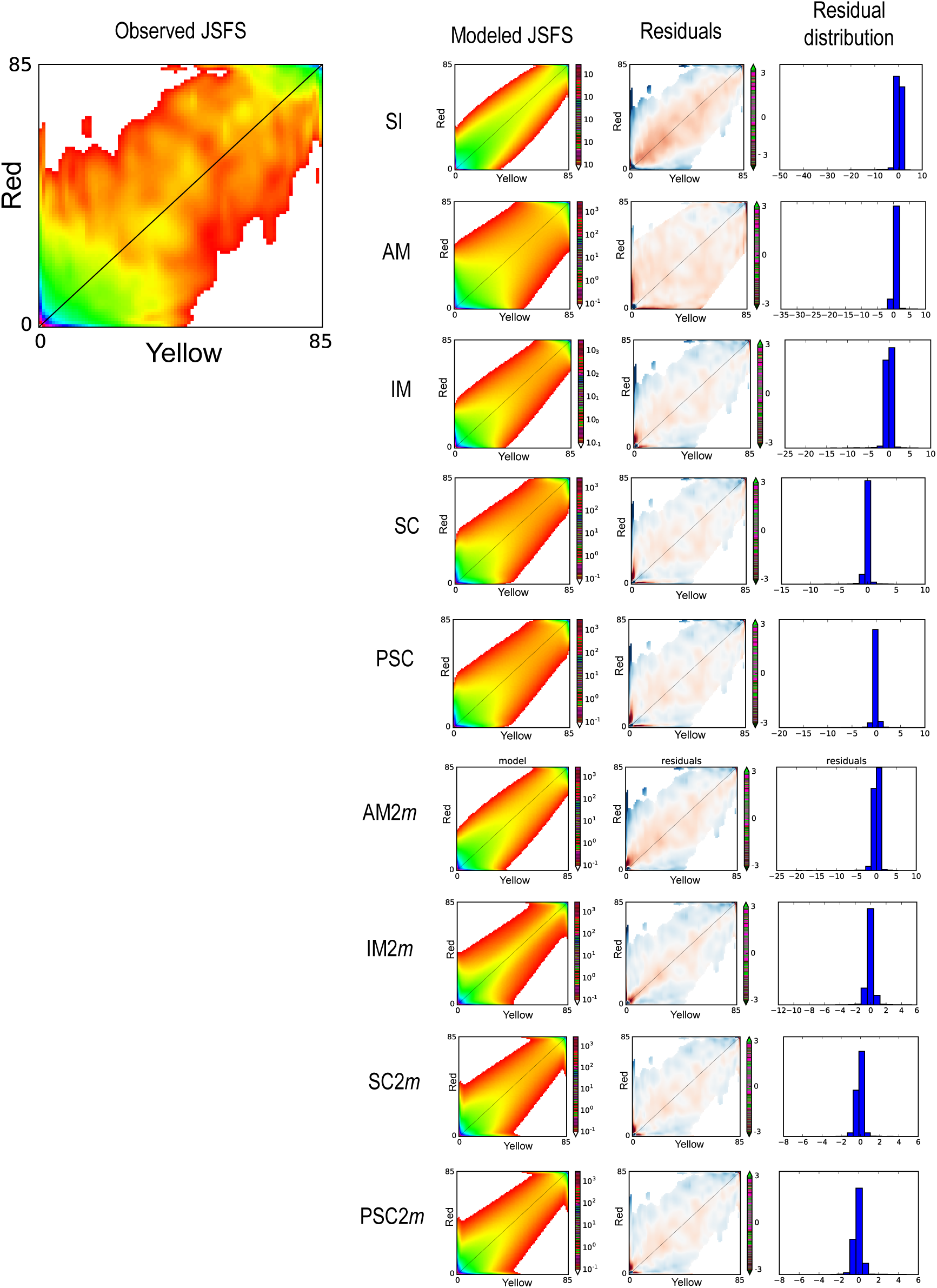
Fits of the different demographic models to the observed data. The observed unfolded joint site-frequency spectrum (JSFS) calculated from the RADseq data set is shown on the left. The first column shows the modeled JSFS for the best fit of each model (See Fig. S1 for a cartoon of each model). The second column shows the residuals of the best model fit to the observed JSFS. The third column shows the residuals plotted as a histogram.

**Figure S5.**
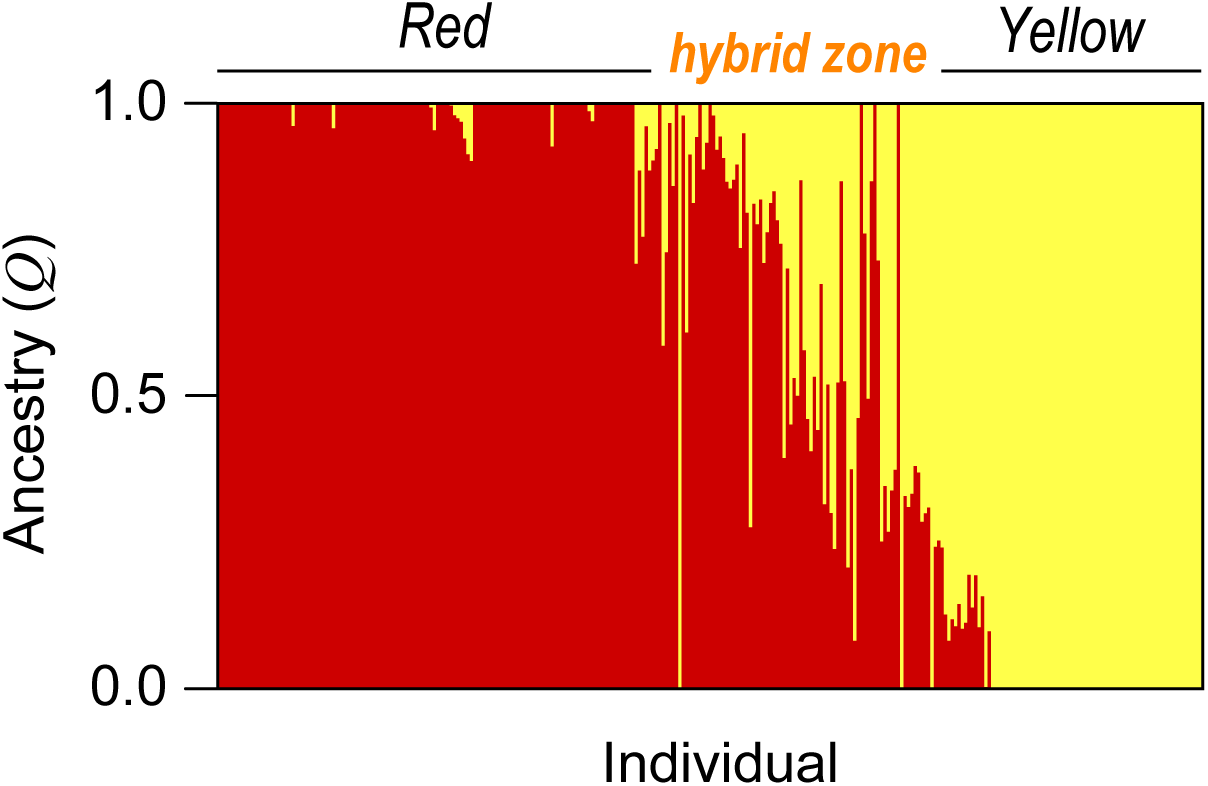
The results of an *Admixture* analysis conducted on the full SNP dataset. Each bar represents an individual and shows its probability of membership (*Q* score) to the two different clusters. Individuals are grouped by whether they come from the distribution of the red ecotype, yellow ecotype, or from within the hybrid zone.

**Figure S6.**
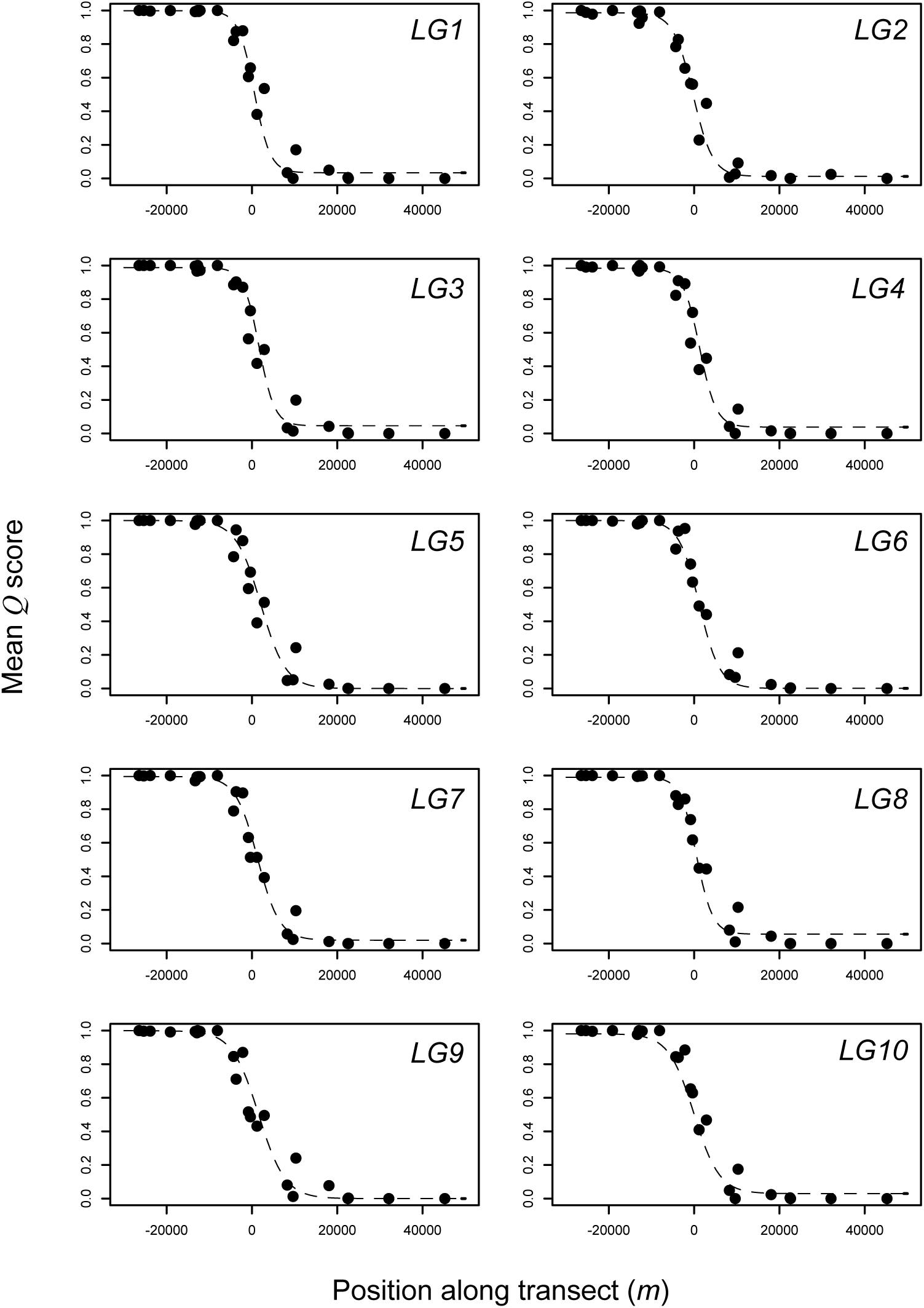
Cline fits for each chromosome. Black points are the mean ancestry scores plotted along the one dimensional transect. The dashed line is the ML sigmoid model.

**Figure S7.**
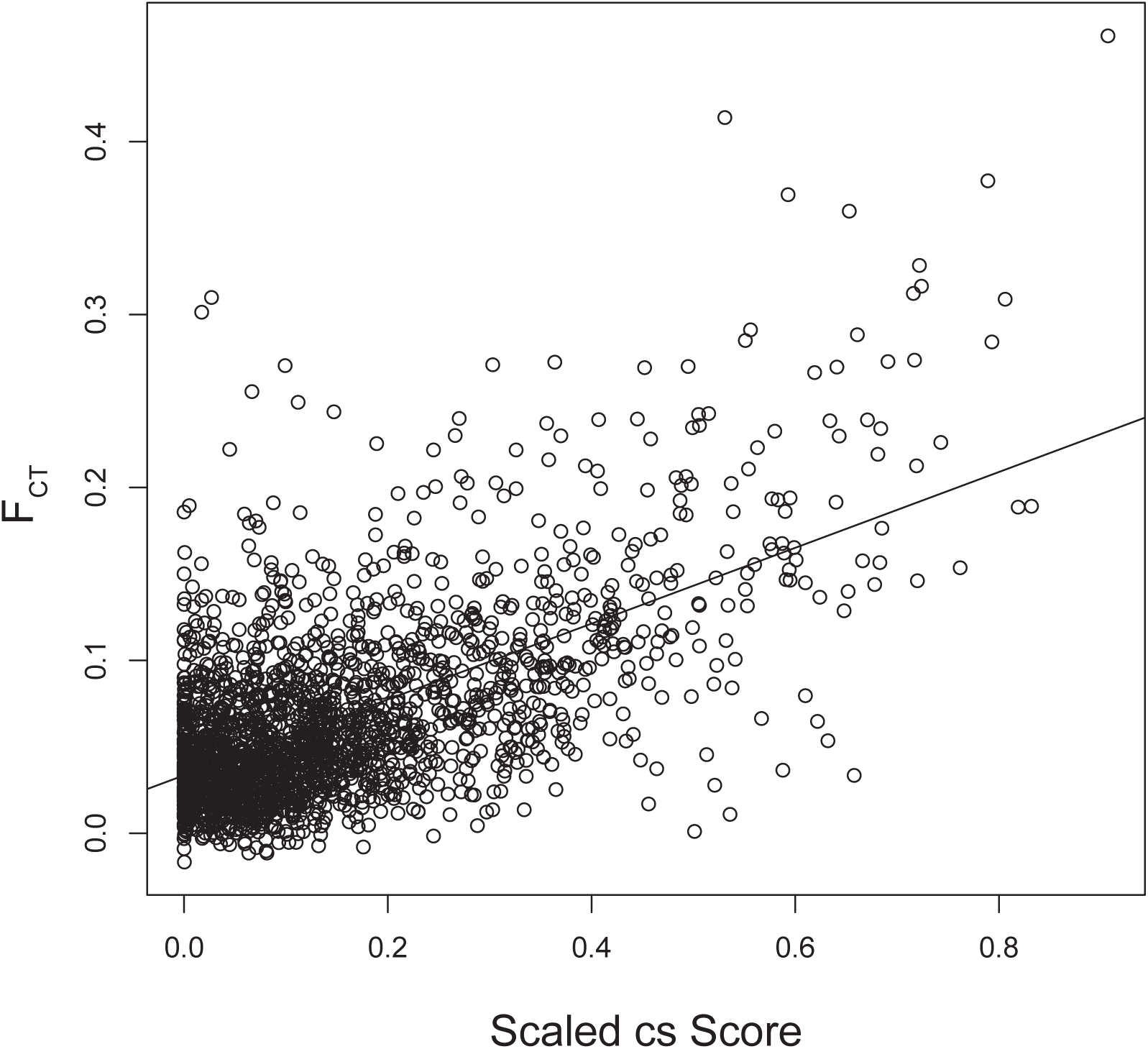
Relationship between the cline similarity score (cs) and *F*_CT_. The line through the plot is the least squares regression (*r*^2^ = 0.38).

**Figure S8.**
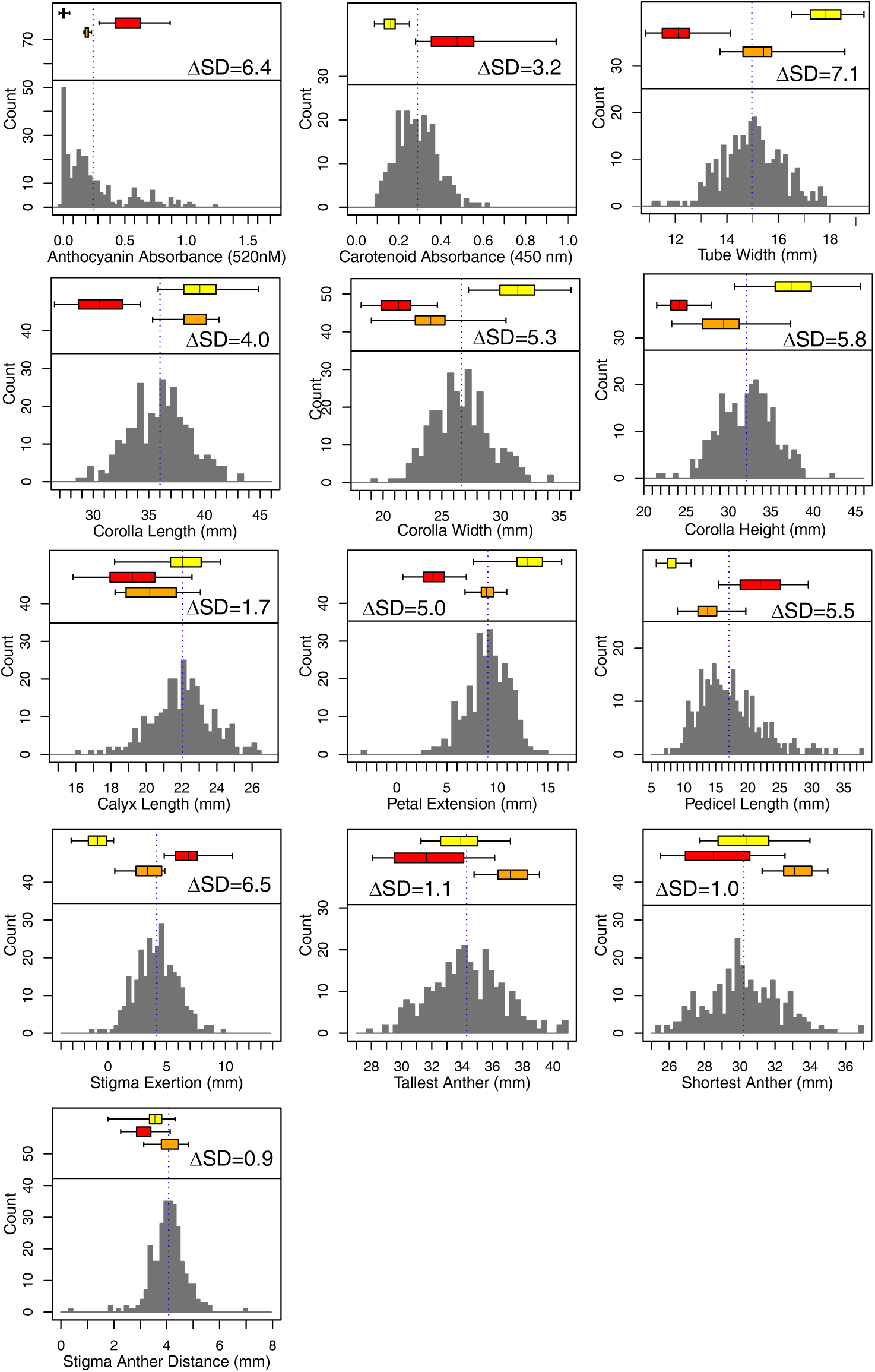
Trait variation in greenhouse-raised red ecotype, yellow ecotype, F_1_, and F_2_ individuals. The histogram in each plot shows the distribution for each trait in the F_2_ population. The top, middle and lower box plots show the distributions for the yellow ecotype, red ecotype and F_1_, respectively. ΔSD indicates the number of standard deviation that the red and yellow ecotypes differ by.

**Figure S9.**
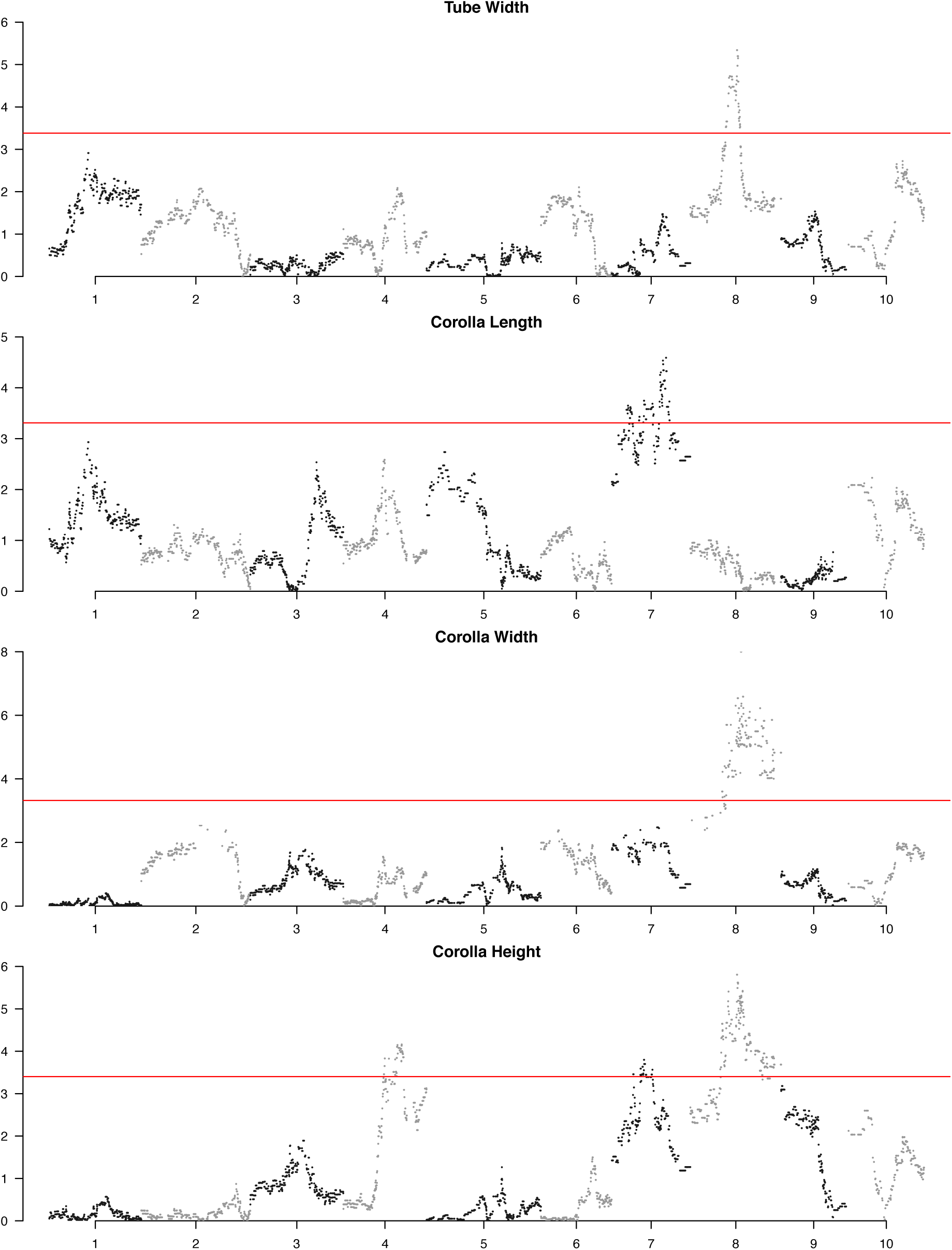
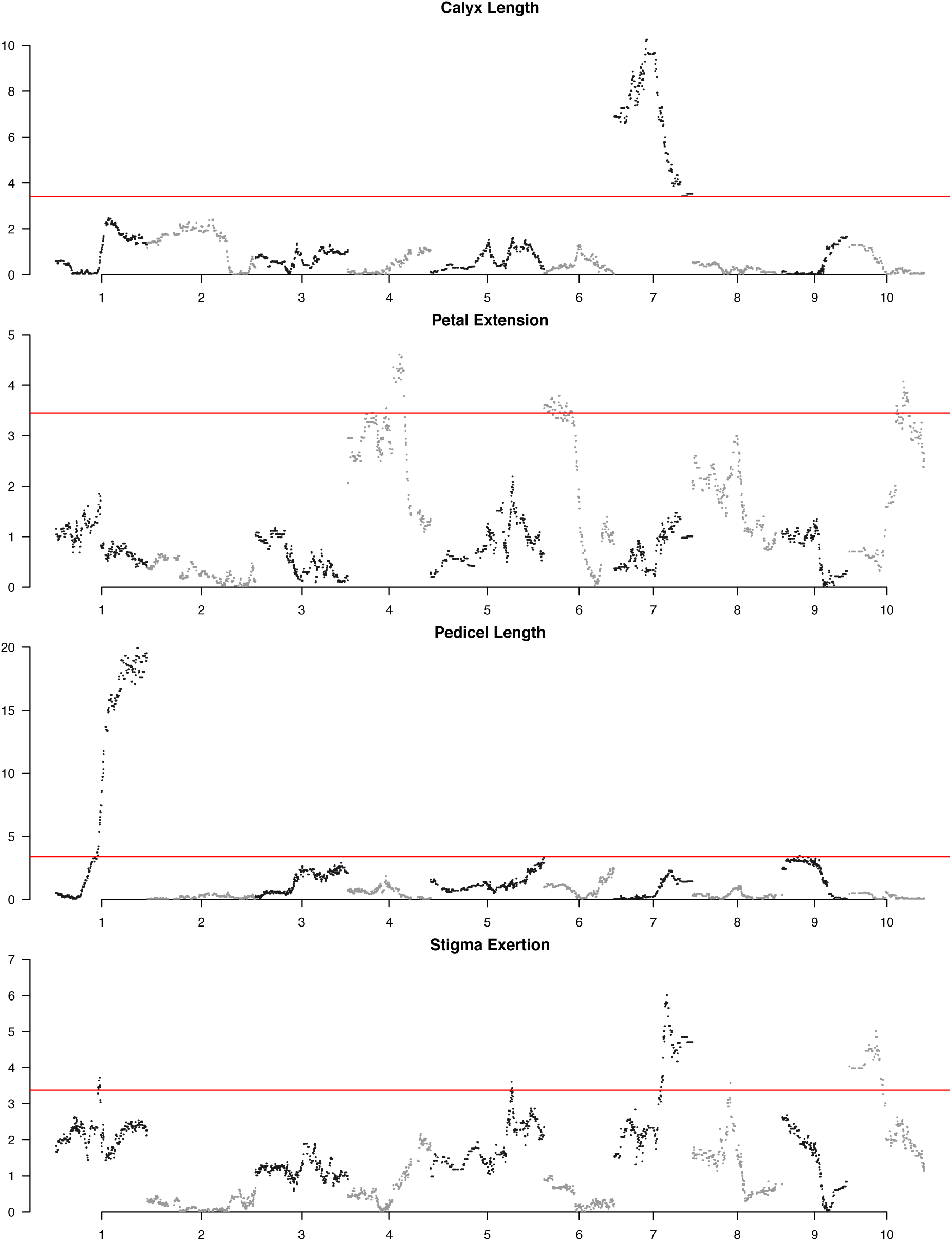
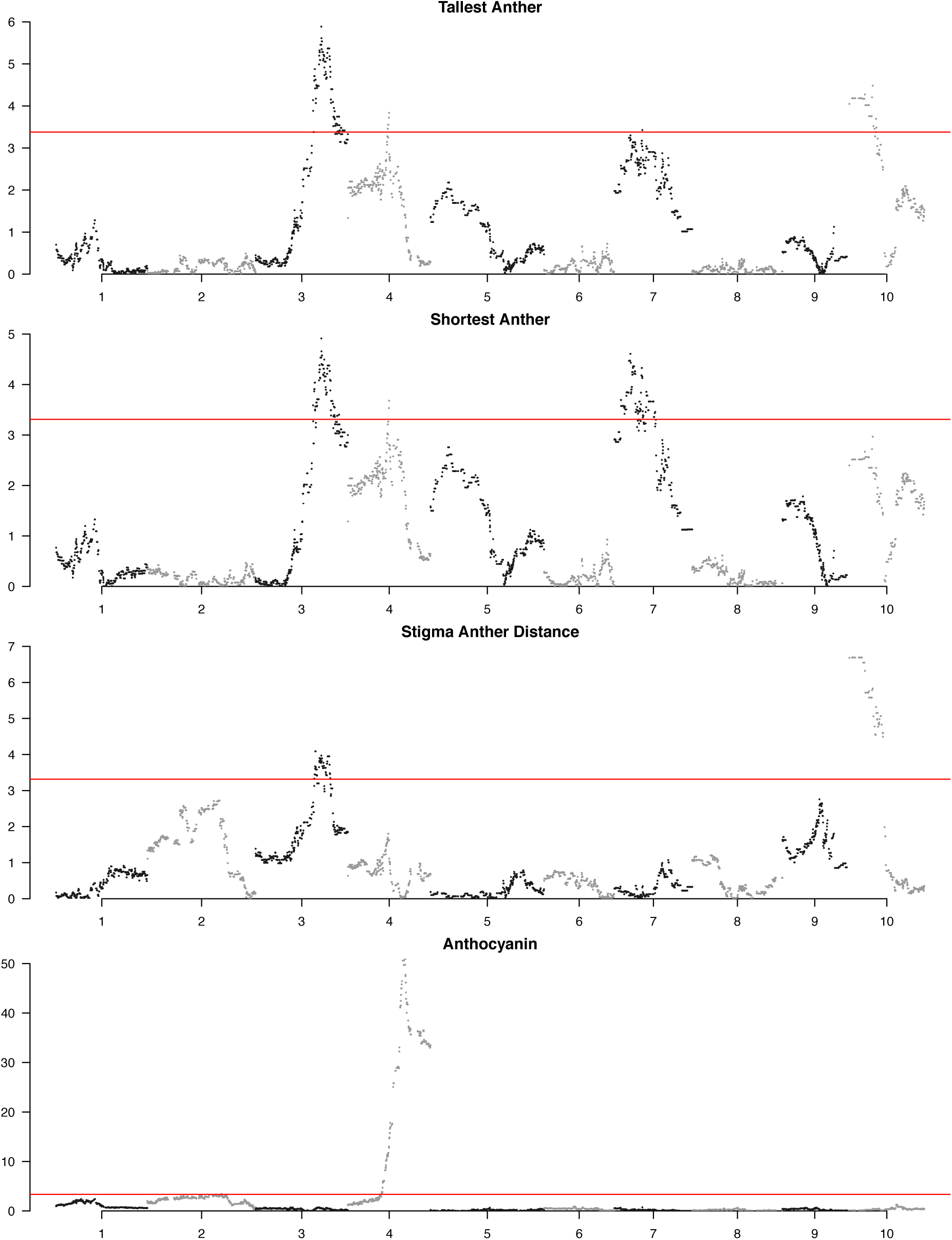
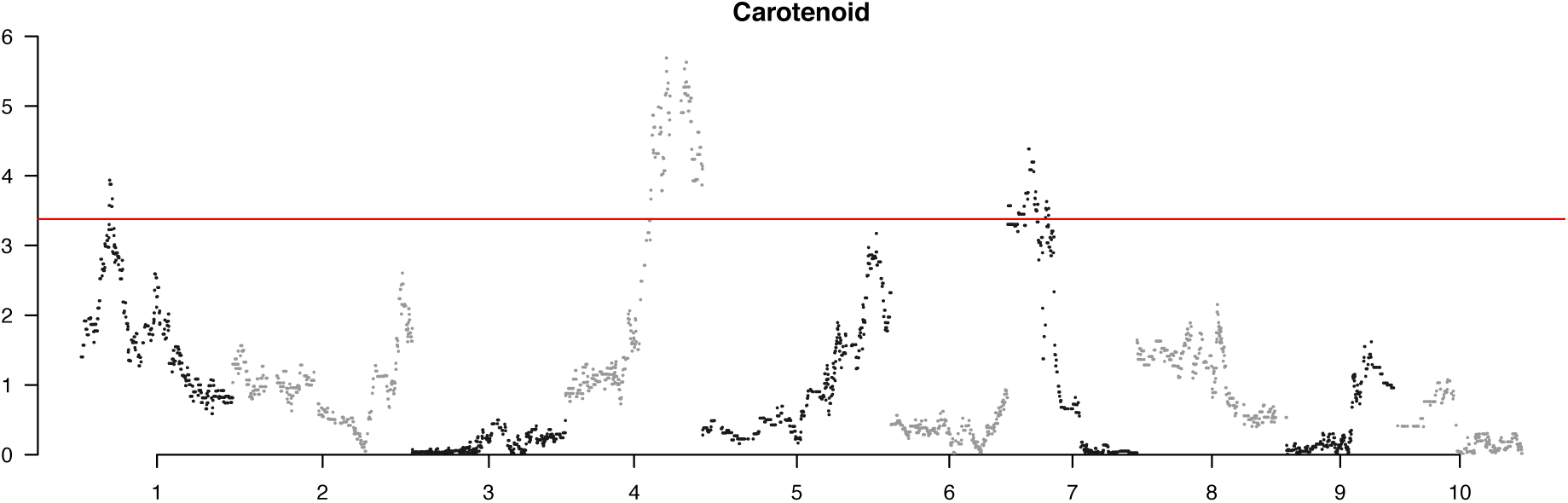
LOD scores plotted across the genome for each trait in the QTL analysis. The horizontal red line shows the LOD score cutoff for the detection of a significant QTL. Note the different y-axis scales among traits.

**Figure S10.**
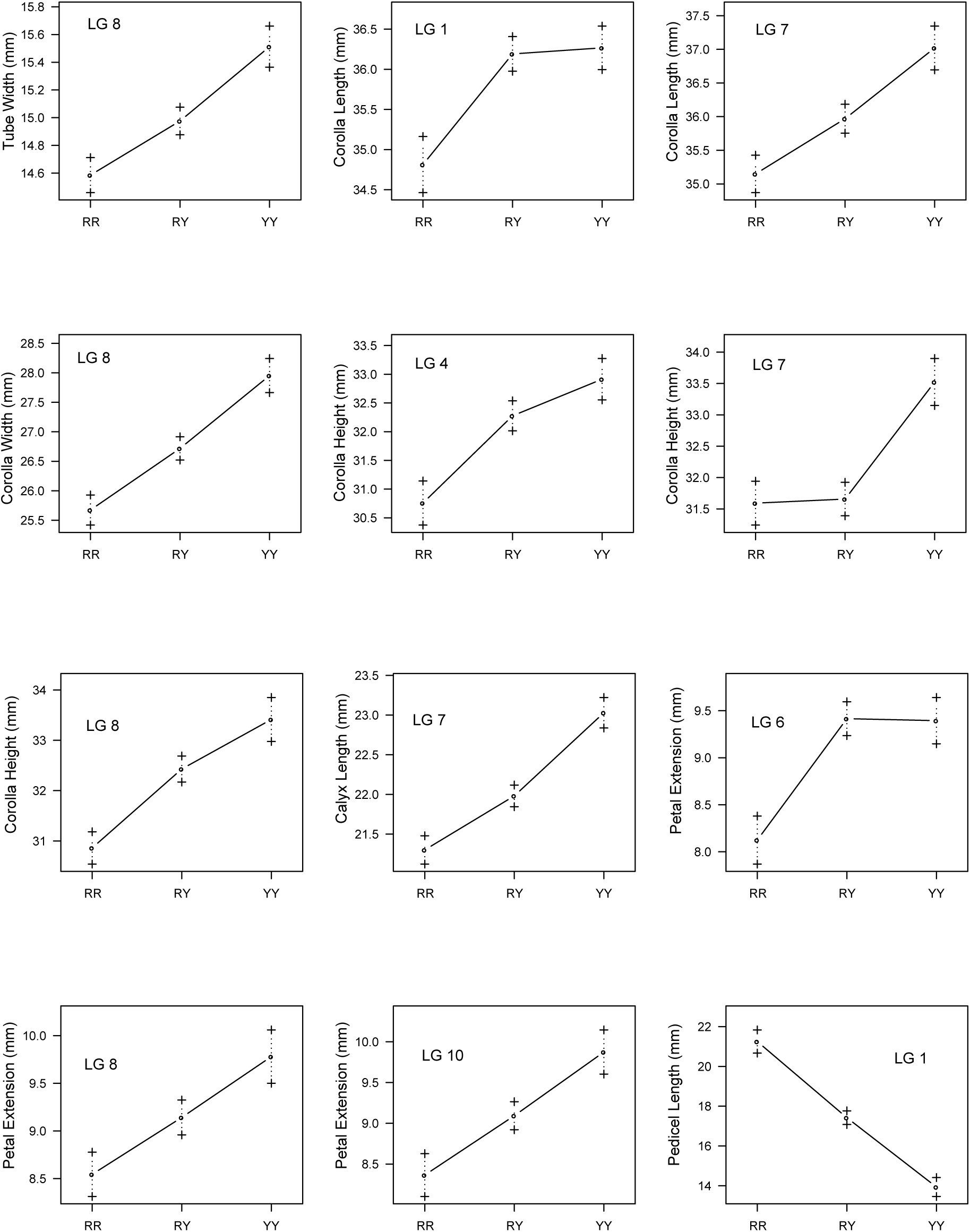
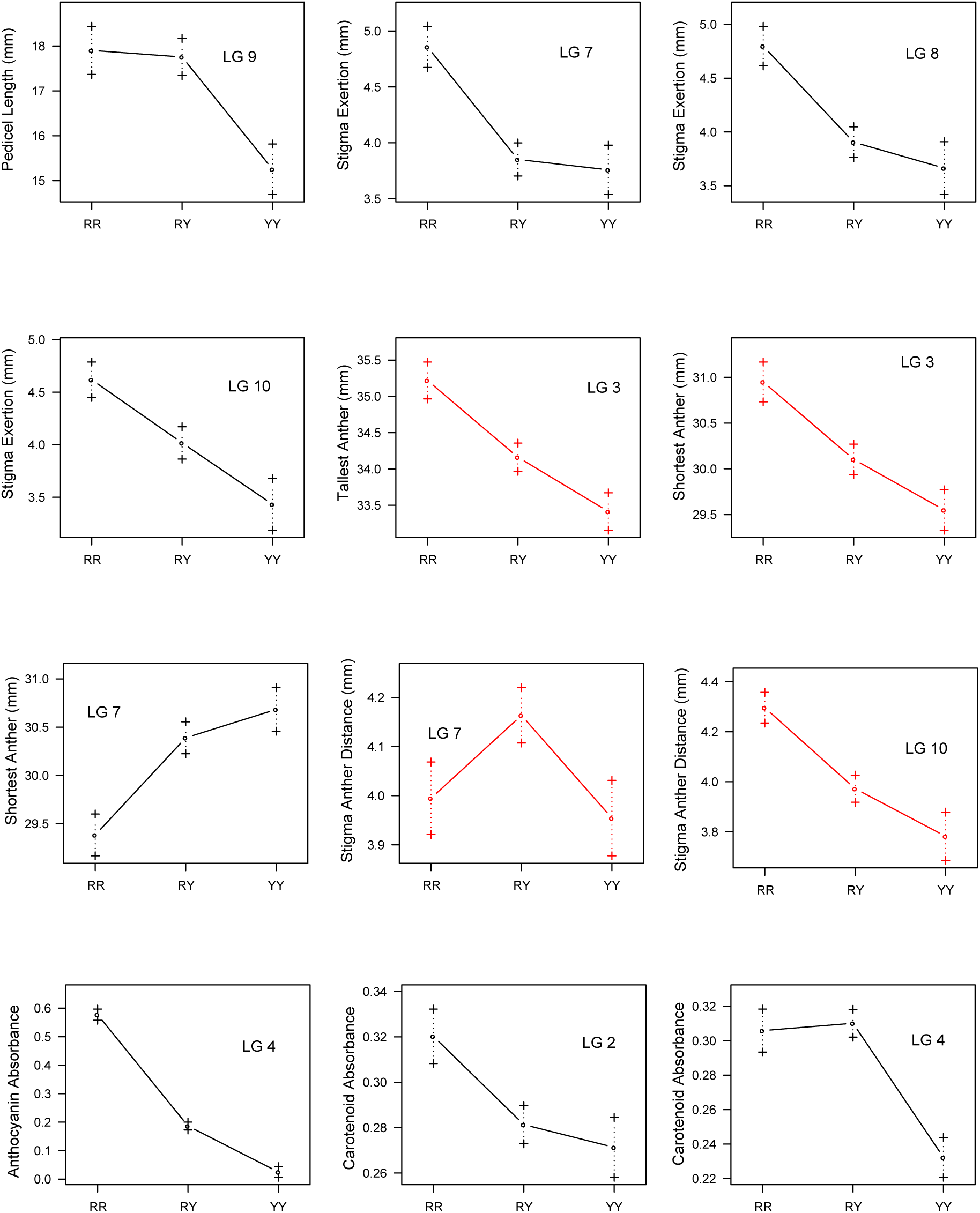
QTL effect plots for each of the 26 QTL discovered in the analysis. Each plot shows the mean (circle) and standard deviation (crosses) for each trait plotted against the three genotypes. RR, both alleles inherited from a red grandparent; YY, both alleles inherited form a yellow grandparent; RY, Heterozygous. Red lines indicate that the effect occurs in the opposite direction of that inferred by the trait differences measured in the red and yellow grandparents (see Fig. S8).

**Figure S11.**
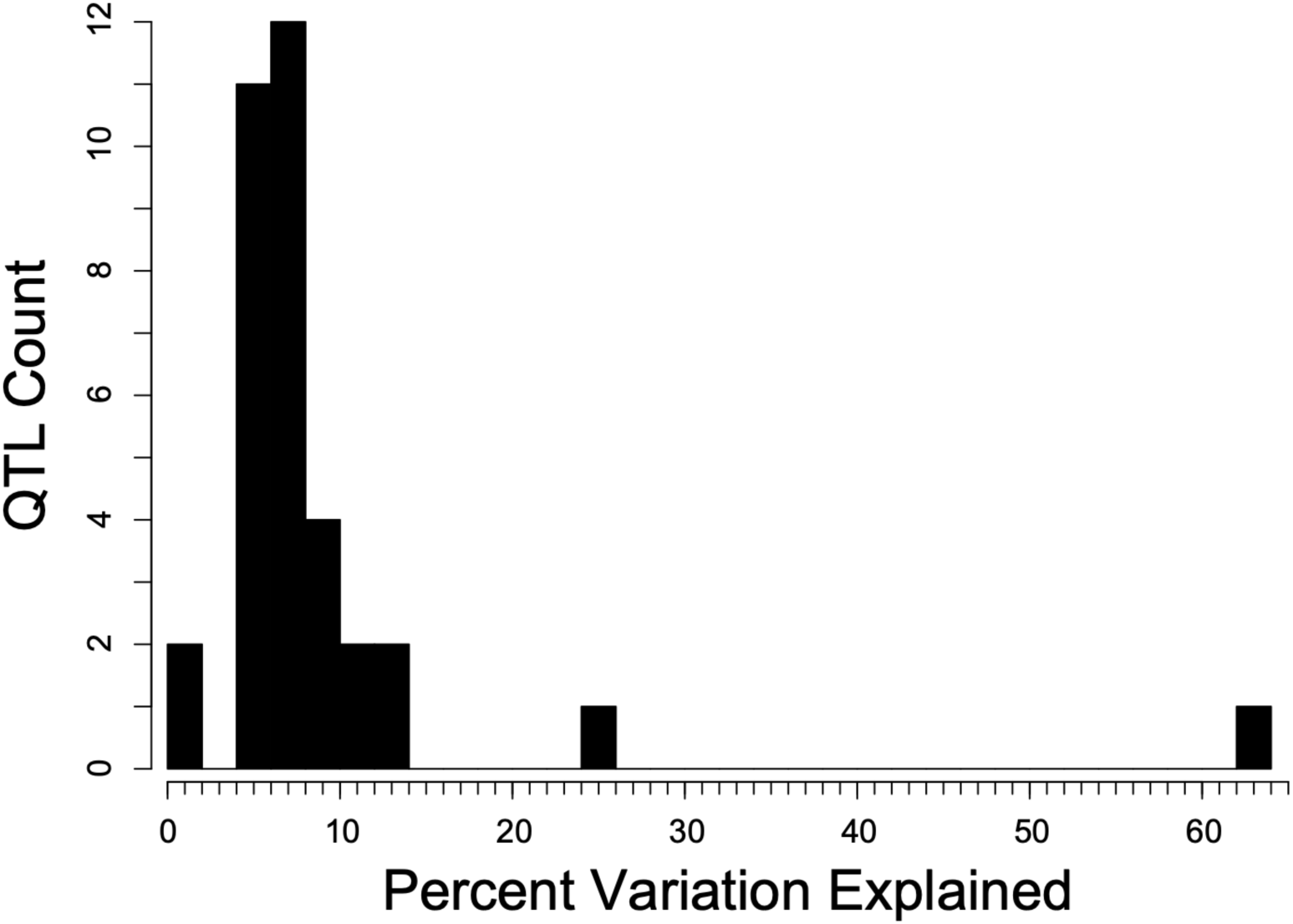
Distribution of effect sizes (percent variation explained among F_2_ plants) for the 26 QTL detected in the study.

**Figure S12.**
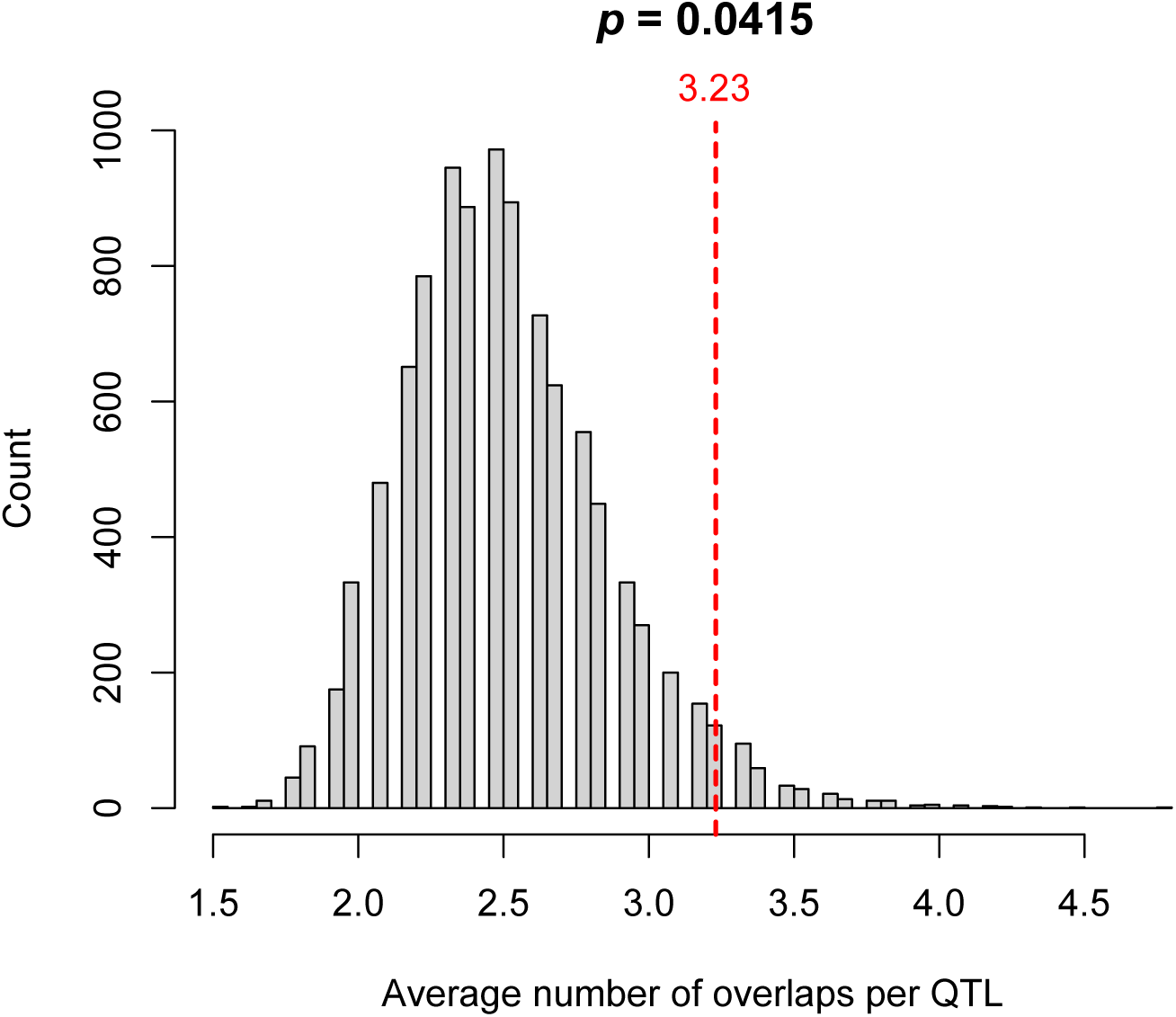
Results from the permutation test for the significant colocalization of QTL. The dashed red line shows the average number of observed overlaps per QTL, while the histogram shows the observed null distribution, generated from 9,999 random permutations. The *p*-value for the test is shown at the top of the plot.

**Figure S13.**
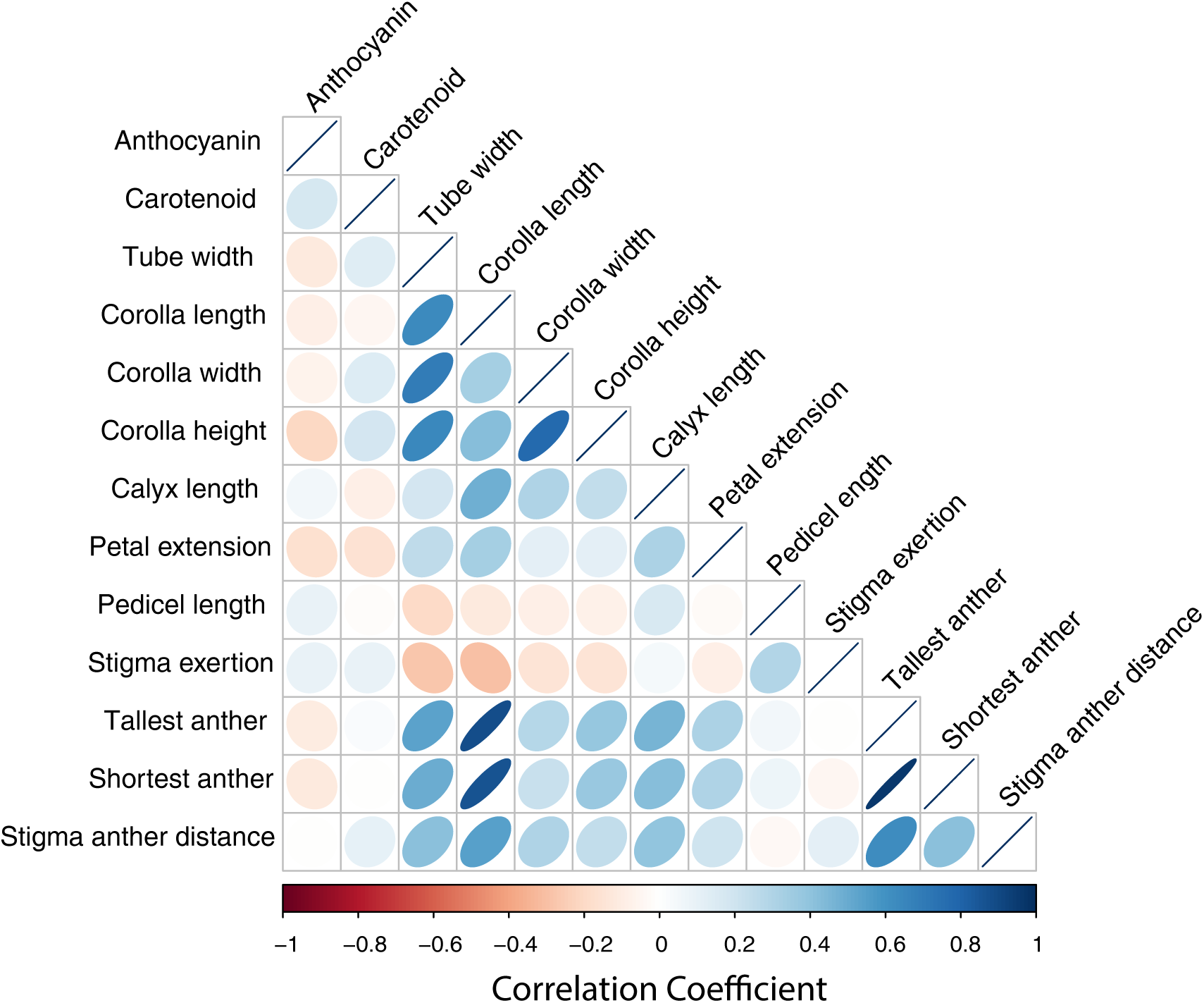
Correlation matrix showing the Pearson correlation between each pair of traits in the F_2_ population. The color and shape of each ellipse indicates the strength and direction of each correlation.

**Figure S14.**
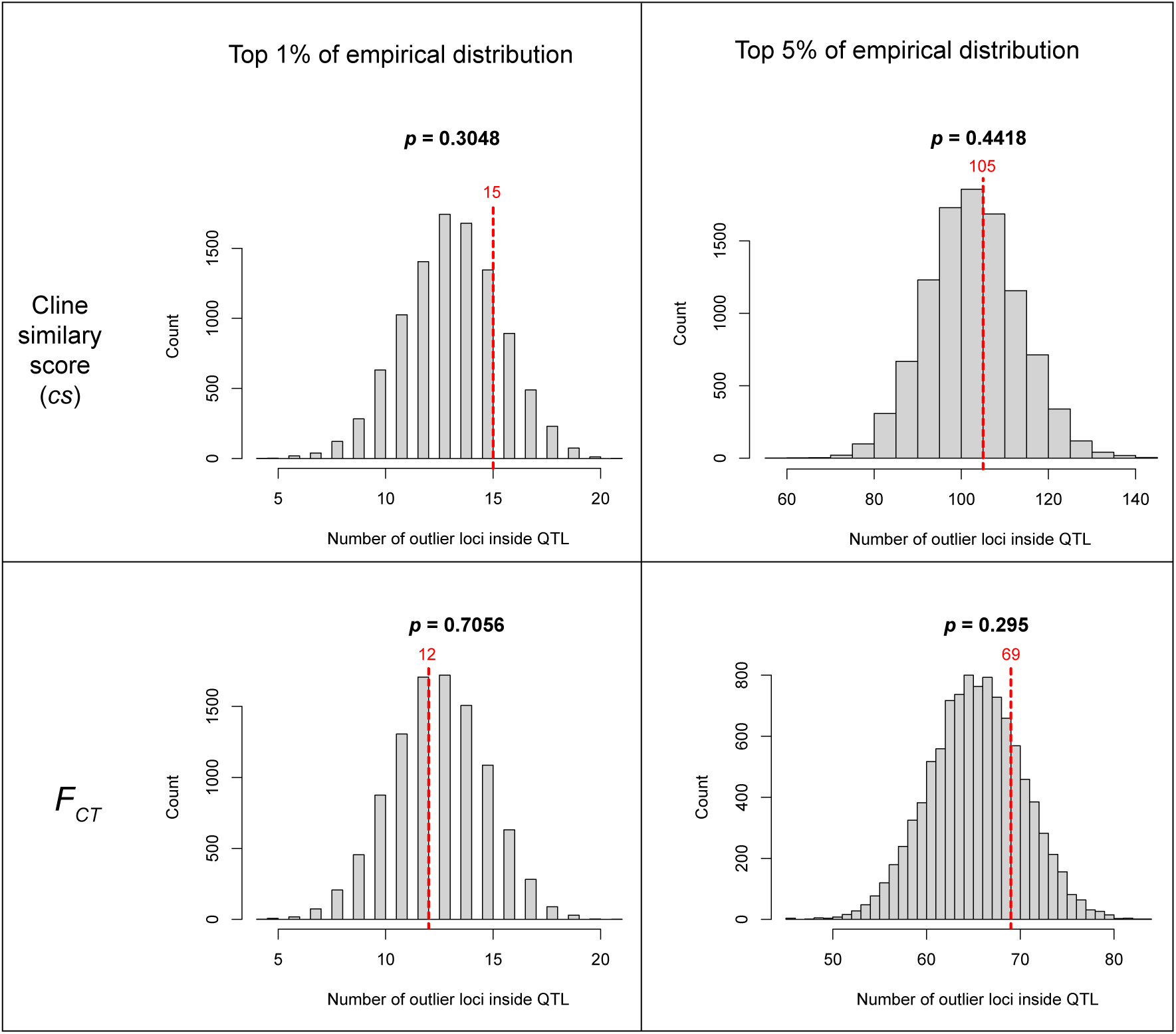
Results from the permutation tests for significant enrichment of candidate barrier loci within the QTL Bayes credible intervals. The test is performed with two different arbitrary cutoffs for defining candidate barrier loci (top 1% and top 5% of the distribution) for two different statistics (the cline similarity score and *F*_CT_). The red line shows the observed number of candidate barrier loci falling within the QTL intervals, while the histogram shows the observed null distribution, generated from 9,999 random permutations. The *p*-value for the test is shown at the top of the plot.

## Supplement 1

### Calculation of ‘cline similarity’ score

#### Rationale

This supplement outlines the rationale behind the calculation of the cline similarity score, which describes the relative shape and position of a cline, relative to some other cline of interest. In our case, we wanted to be able to view the variation in multiple parameters that describe different features of a geographic cline in an integrated way, so we could do things, like plot clinal variation along a genome.

Consider a simple, sigmoid cline, like the one shown in Figure A. This cline model, which is the most commonly used in empirical studies, is described by 4 fitted parameters: the cline centre (*c*), cline width (*w*), the mean trait value on the ‘high’ side of the cline (*Q*_max_), and the mean trait value on the ‘low’ side of the cline (*Q*_min_). The last 2 parameters can be reduced to a single parameter, Δ*Q* = *Q*_max_ - *Q*_min_, which quantifies the total change in a trait across the transect. It is worth noting that we use the notation Δ*Q* to denote change in ancestry scores across the cline, rather than Δ*z*, which is typically used for a quantitative trait (or Δ*p* for allele frequency).

**Fig. A.**
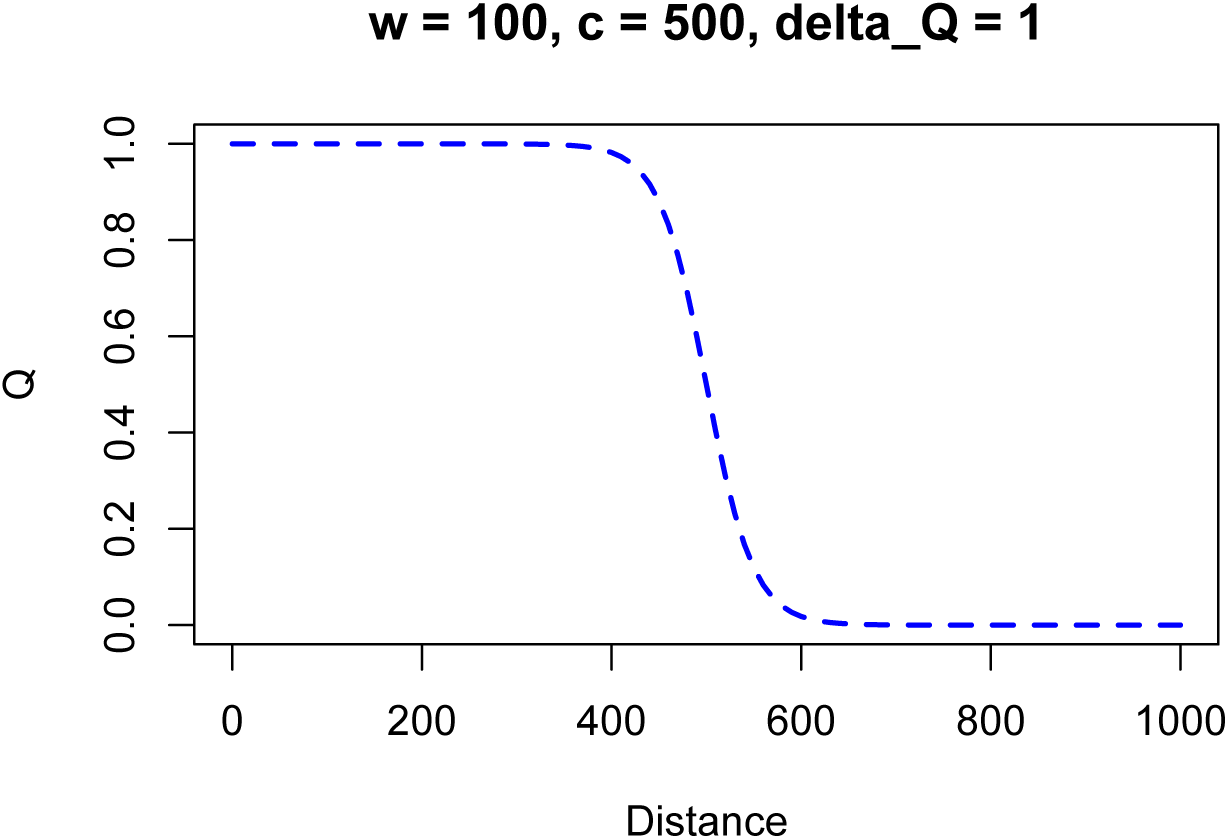
Simple sigmoid cline model described by 3 parameters, Δ*Q*, *w*, and *c*.

Each parameter tells us something about the feature of the cline and is relevant to inferences that we might make about selection in a hybrid zone (provided that we are happy to make some assumptions). For example, the width of the cline is inversely proportional to the strength of selection acting on the trait; the centre tells us about the spatial pattern of selection across the hybrid zone; and the change in the trait value across the cline tells us about the strength of the difference in the trait across the cline. In a situation where selected and neutral loci are at equilibrium, the above cline would indicate that variation in the trait is maintained by selection.

The problem is that inferences cannot easily be made from a single cline parameter. This is highlighted in Figure B, where several clines are shown, with certain parameters fixed and others varying. The left plot shows three clines with the same values of *w* and *c*, but substantially different values of Δz; The middle plot shows multiple clines with the same values of Δz and c, but with different values of *w*; the third shows clines with identical shapes (i.e., same Δ*z* and *w*), but different values of *c*.

**Fig. B.**
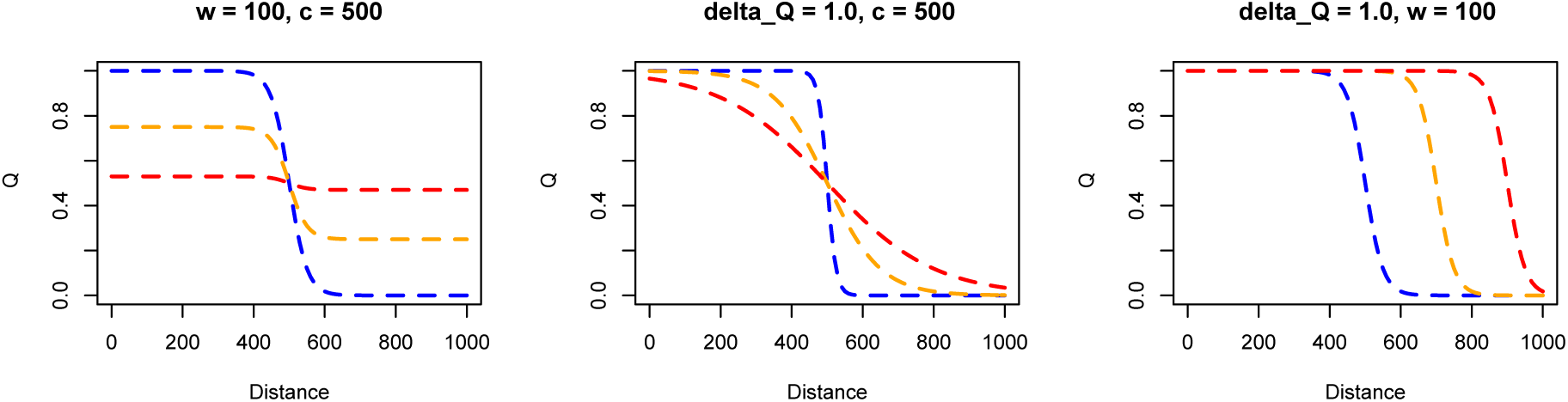
**Examples of clines where a single parameter varies but the other two parameters are fixed.**

In the above plots, we can easily evaluate each cline in terms of its fit to some hypothesis by visually inspecting them. However, when we have many clines, this is impractical and highly subjective. But these plots clearly show that we cannot infer much by looking at the distribution of a single parameter, especially in real data where all parameters will vary simultaneously.

#### The ‘cline similarity’ score

To help simplify the process of identifying clines of interest, we created an *ad-hoc* statistic that we call the ‘cline similarity’ or ‘*cs*’ score. The *cs* score summarises cline shape and relative position with a single number that can be calculated from the ML cline parameters. *cs* is calculated as:

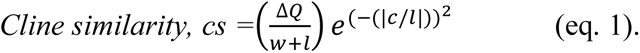

Where Δ*Q w* and *c*, are defined as above and *l* is a scaling factor that will be explained further below. Eq. 1 can be broken down into two terms that determine the *cs* score in different ways. The first term is a shape score, that describes the shape of a cline, independent of the location of the cline centre:

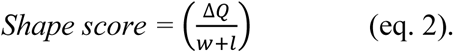

The second term is a centre penalisation score, which downgrades the shape score based on the value of the fitted cline centre relative to some point of reference (explained further below).

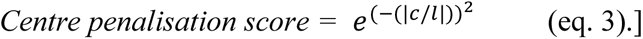

These formulae and the decisions made to arrive at them will next be explained by walking through its calculation for the *Mimulus* dataset.

#### Application to the *Mimulus* dataset

For the *Mimulus* dataset, we fit clines to ancestry scores in 2,173 non-overlapping windows, each containing 100 SNPs. The details of how the fits were done are described in the main text, but the clines can be fitted in any way. We only need the parameters for the fitted model to calculate *cs*.

Starting with the shape score, we wanted a quantity that described variation in the shape of the cline. Ultimately, we wanted clines with a small *w* and large Δ*Q* to have high shape scores, and clines with very low Δ*Q* and high *w* to have a low shape scores. Figure C shows the joint distribution of Δ*Q* and *w* for all of the windows. The numbers in the plot correspond to the cline fits for 10 selected windows, which are shown on the right side of the plot. The orange star (cline 2), is not for a specific window, but rather is the fit to mean *Q* score for each location, averaging across all windows (i.e., it is a cline fitted to the mean of means for each location). The green star shows the location of the genome-wide cline in this space (i.e., the one shown in figure 3a of the main text, and figure D (*ii*)).

**Fig. C.**
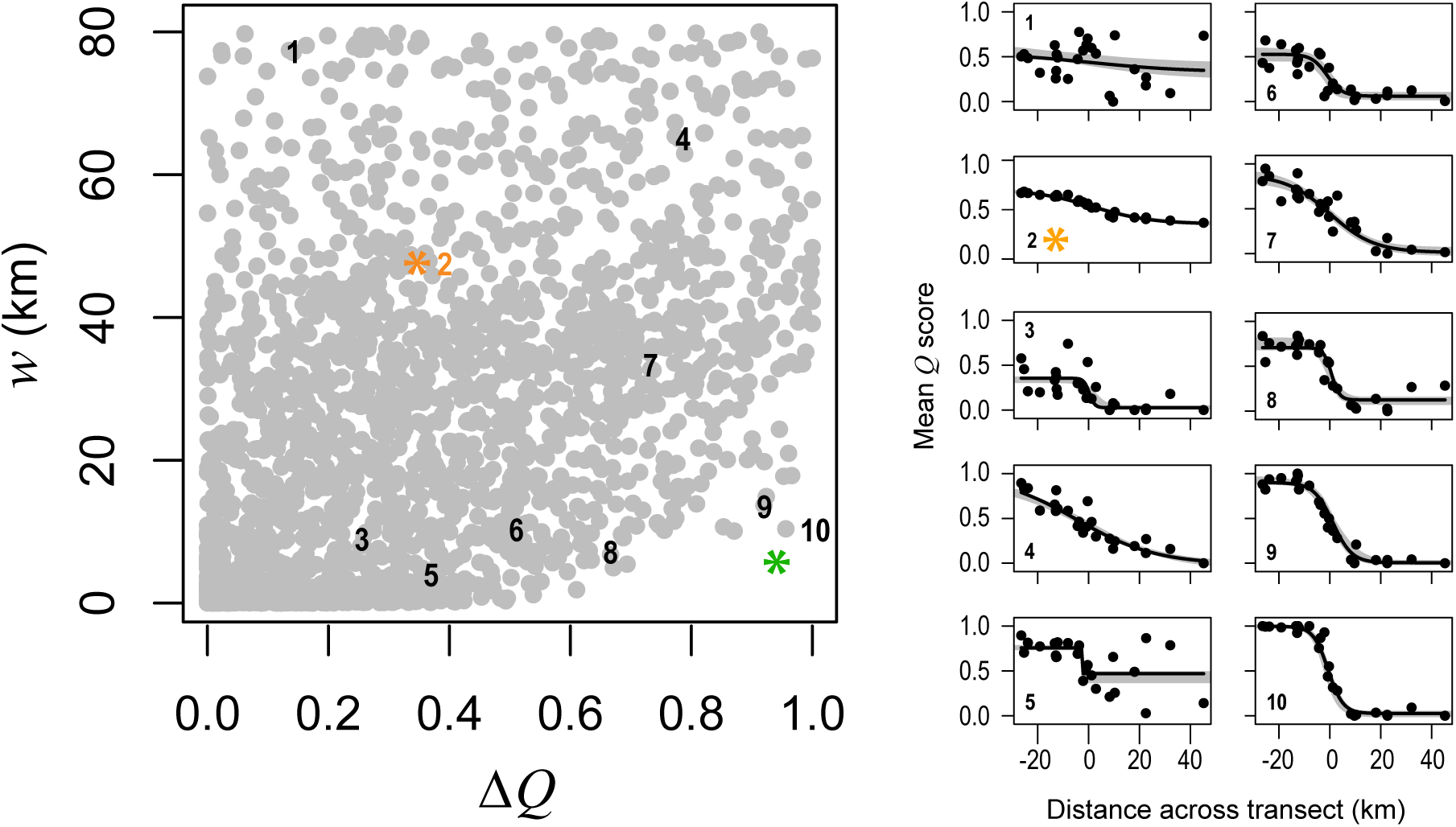
Joint distribution of *w* and Δ*z* for the *Mimulus* dataset. The clines on the right show fits that correspond to the numbers in the bivariate plot.

The bottom right corner of Figure C contains the clines that are most interesting in terms of our hypothesis regarding barrier loci. The ones in the top left and bottom left are of little interest, because Δ*Q* is always very small. The ones in the top right are more interesting, because they show high Δ*Q* but are very wide; these clines could be generated by processes like isolation by distance, or the collapse of secondary clines when barriers to gene flow are porous.

Figure D*(i)* shows the same plot, but with the points coloured by the shape score (10 coloured bins, but the values are continuous), as given in Eq. 2. Here we define *l*, the scaling parameter, as 0.5*t*, where *t* is the length of the transect. *l* determines precisely how variation in the shape scores is spread out across the joint distribution of Δ*Q* and *w*. In our case, 0.5*t*, gives a distribution shape scores that suits our purposes, but other values might be more suitable for other study systems.

**Fig. D.**
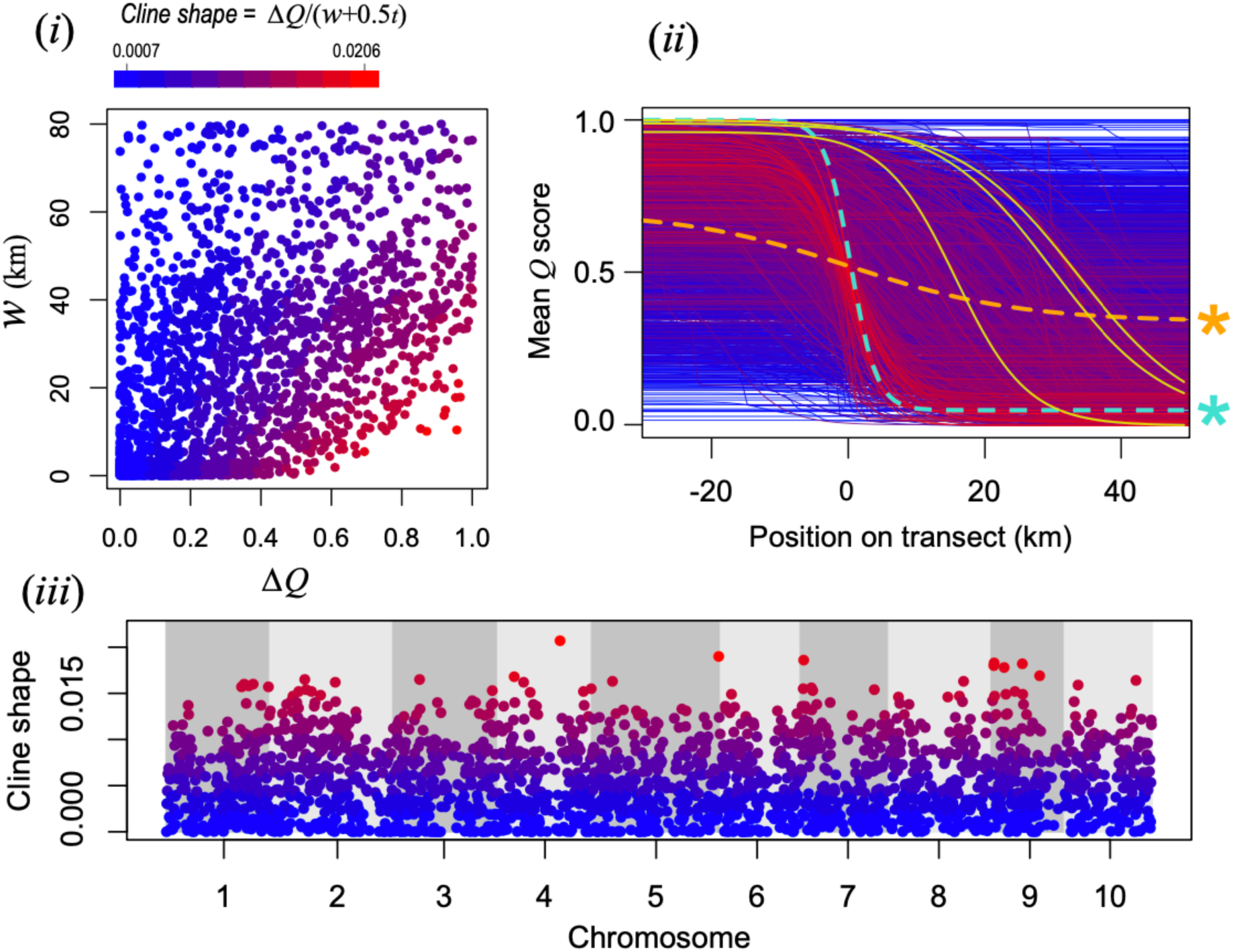
Shape scores for the 2,173 windows. (*i*) Joint distribution of *w* and Δ*Q*, coloured by the shape score. (*ii*) Cline models coloured by the shape score. (*iii*) The shape scores plotted across the genome. See text for more details.

By colouring the cline models for each window by their shape scores, as shown in Figure D(*ii*), we can see that clines with ‘hotter’ colours do have shapes more similar to the genome wide cline (the dashed cyan line) than ‘cooler’ colours. However, it is also clear that some clines with high shape scores have centres that are displaced a very long way from the genome wide cline. Some of these have been coloured solid yellow to help highlight them.

Although these clines are interesting for a variety of reasons, they are not spatially associated with the centre of the hybrid zone that also coincides with the transition in floral trait differences. Ideally, we would like our summary statistic to also reflect where the cline is positioned in relation to this location. To do this, we penalise the shape score based on the position of the cline centre, relative to a position of interest, which needs to be given position 0. This could be a feature of the environment or a cline in a focal marker or trait. In our case, the position of interest is the centre of the genome-wide ancestry cline.

Figure E*(i)* shows how the value of the inferred centre informs the centre penalisation function outlined in Eq. 3. If the fitted cline centre coincides exactly with the point of interest (*c* = 0), the *cs* score for that cline is simply the shape score (i.e., *cs* = shape score). However, as the difference between fitted centre and point of interests increases (i.e., |0 - c|), then the shape score is downgraded according to the centre penalization function, resulting in a *cs* score lower than the shape score.

**Figure E.**
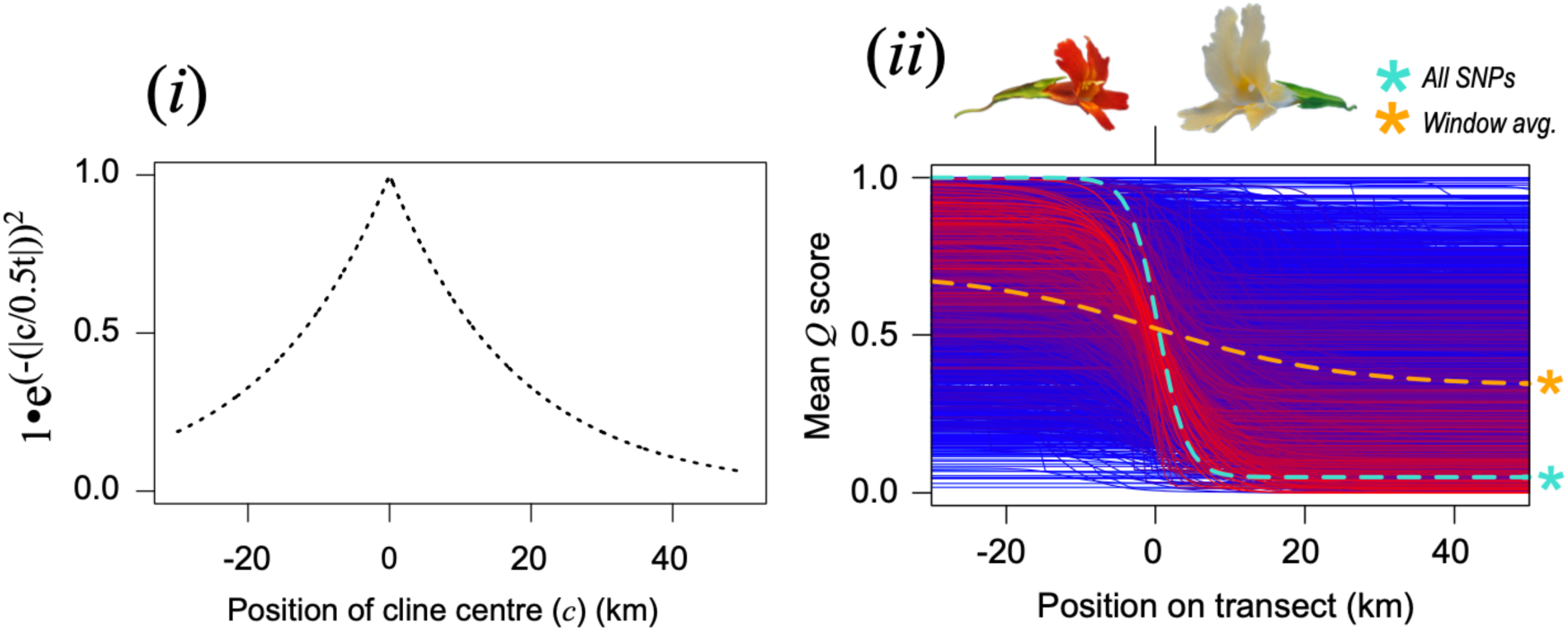
Penalisation of the shape score to give the final *cs* score. (*i*) The effect of the cline centre penalisation function on the shape score. Here, an arbitrary shape score of 1 is used for illustrative purposes. (*ii*) The cline models for all windows after the cline penalisation score, coloured by the overall *cs* score.

The results of this final transformation step can be seen in figure 3c of the main text, which is reproduced in Figure E(*ii*) for convenience. In that figure, the *cs* scores are scaled between 0 and 1, where 0 is the cline with the lowest *cs* score, and 1 is the *cs* score for the genome-wide cline.

#### Some afterthoughts (if others attempt something similar)

This was done 5 years ago and if I was doing this again from scratch, I would probably think about it a bit differently. First, I think it might be useful to try and incorporate the variance explained by the cline model into the *cs* score. i.e., if the data around the cline are extremely noisy (common for clines with a small Δ*Q*, e.g., cline 1 in Figure C ), we might conclude that these are less likely to be of interest. The variance explained was used primarily in Westram *et al.* 2017 and seemed to be good at identifying non-neutral clines in simulations.

